# A two-component quasi-icosahedral protein nanocompartment with variable shell composition and irregular tiling

**DOI:** 10.1101/2024.04.25.591138

**Authors:** Cassandra A. Dutcher, Michael P. Andreas, Tobias W. Giessen

**Affiliations:** Department of Biological Chemistry, University of Michigan Medical School, Ann Arbor, MI 48109, USA

## Abstract

Protein shells or capsids are a widespread form of compartmentalization in nature. Viruses use protein capsids to protect and transport their genomes while many cellular organisms use protein shells for varied metabolic purposes. These protein-based compartments often exhibit icosahedral symmetry and consist of a small number of structural components with defined roles. Encapsulins are a prevalent protein-based compartmentalization strategy in prokaryotes. All encapsulins studied thus far consist of a single shell protein that adopts the viral HK97-fold. Here, we report the characterization of a Family 2B two-component encapsulin from *Streptomyces lydicus*. We show the differential assembly behavior of the two shell components and demonstrate their ability to co-assemble into mixed shells with variable shell composition. We determined the structures of both shell proteins using cryo-electron microscopy. Using 3D-classification and crosslinking studies, we highlight the irregular tiling of mixed shells. Our work expands the known assembly modes of HK97-fold proteins and lays the foundation for future functional and engineering studies on two-component encapsulins.

## Introduction

The HK97 fold is principally found in the major capsid proteins of bacteriophages of the ubiquitous viral order Caudovirales (*1*). It is named after the fold observed in the major capsid protein of Escherichia virus HK97, a double-stranded DNA bacteriophage natively infecting *Escherichia coli* (*2*). The HK97 fold is also found as a key structural component in other icosahedral protein assemblies including tailed archaeal viruses (*3*), the floor domains of animal-infecting viruses of the order Herpesviridae (*4*), and the shell proteins of encapsulins – self-assembling enzyme-loaded protein nanocompartments – found in many bacteria and archaea (*5–7*).

The HK97 fold is characterized by three highly conserved structural features, the axial domain (A-domain), peripheral domain (P-domain), and the extended loop (E-loop) (Fig. 1A) (*2, 7*). The structural plasticity and versatility of the HK97 fold is highlighted by the fact that its three core structural features – A-domain, P-domain, and E-loop – can be maintained in proteins with little sequence similarity while a range of diverse insertion domains have been found embedded at various positions within the HK97 fold (*8, 9*). In addition, icosahedral HK97 assemblies have been shown to exhibit dynamic pores able to open or close depending on the particular stimuli (*10, 11*).

**Figure 1.**
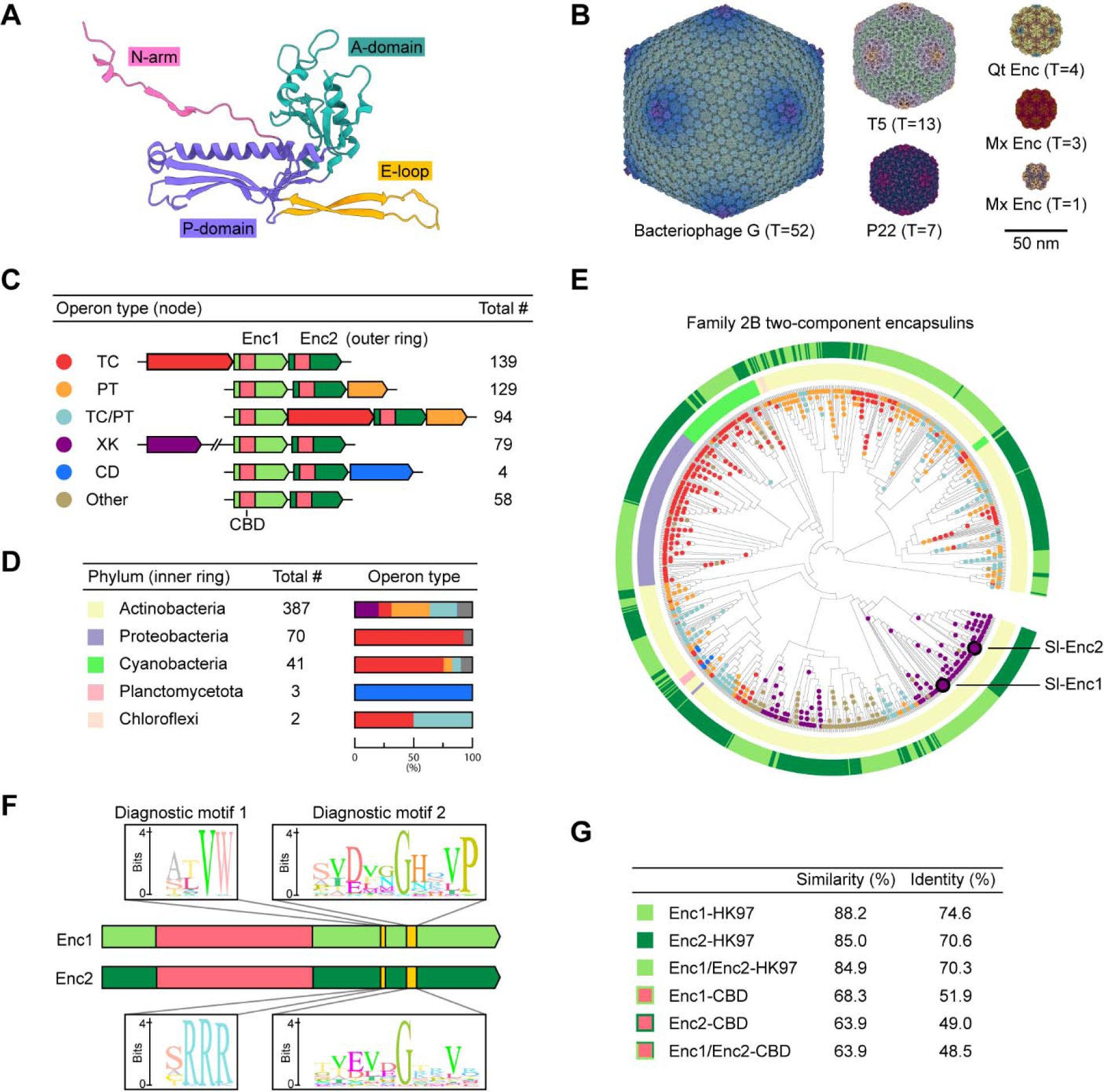
The HK97 fold and distribution and diversity of Family 2B two-component encapsulin systems. (**A**) HK97 bacteriophage protomer (chain A) highlighting the canonical HK97-fold (PDB ID: 1OHG). N-arm: N-terminal arm. A-domain: axial domain. E-loop: extended loop. P-domain: peripheral domain. (**B**) HK97-fold viral capsids and encapsulin shells exhibiting varying triangulation (T) numbers. Radially colored cryo-EM maps are shown. Bacteriophage G (EMD-8484), T5 (EMD-20122), P22 (EMD-35120), *Quasibacillus thermotolerans* (Qt) encapsulin (EMD-9383), *Myxococcus xanthus* (Mx) encapsulin (T=3: EMD-41322, T=1: EMD-24815). (**C**) Two-component Family 2B encapsulin operon types. TC: terpene cyclase, PT: polyprenyltransferase, XK: xylulose kinase, CD: cysteine desulfurase, CBD: cyclic nucleotide (cNMP)-binding domain. (**D**) Phylogenetic distribution of two-component encapsulin systems. **(E)** Phylogenetic tree of all identified shell proteins encoded in two-component Family 2B encapsulin operons. Outer ring: Enc1 vs Enc2. Inner ring: phylum. Nodes colored by cargo type. **(F)** Domain organization of the two *Streptomyces lydicus* (Sl) shell proteins highlighting two conserved sequence differences found in all Enc1 and Enc2 shell proteins (diagnostic motif 1 and 2). (**G**) Comparative sequence analysis of Enc1 and Enc2 shell proteins. HK97: HK97-domain.

Protein shells (encapsulins) and capsids (viruses) that utilize the HK97-like fold exhibit an impressively large size distribution, ranging from 18 nm (triangulation number (T) of T=1, 60 protomers; empty *Myxococcus xanthus* encapsulin) to 180 nm (T=52, 3120 protomers; bacteriophage G) in diameter (Fig. 1B) (*12, 13*). To assemble icosahedra greater than T=1, both pentamers and hexamers are needed to tile a closed shell or capsid (*14, 15*). Nearly all characterized HK97 fold protein assemblies use a single major capsid or shell protein to form icosahedral structures, relying on the built-in conformational flexibility of the HK97 fold, especially the E-loop, to assemble both oligomeric states – pentamers and hexamers – needed for assembling closed shells or capsids. There is a single virus – bacteriophage T4 – that utilizes two distinct HK97 fold proteins for separate structural roles, forming either homo-pentamers or homo-hexamers in the context of its assembled capsid (*16, 17*).

Encapsulins are the only non-viral structures known to utilize the HK97 fold (*5–7*). All so far studied encapsulins are composed of a single type of protomer that self-assembles into defined shells with sizes ranging from 18 to 42 nm in diameter and triangulation numbers of T=1, T=3, or T=4 (*7, 18, 19*). Many prokaryotes utilize encapsulin shells to compartmentalize specialized metabolic processes (*20, 21*). This is accomplished through the selective encapsulation of specific enzymes, targeted to the encapsulin lumen via a C- or N-terminal targeting peptide or domain present in all native cargo proteins (*22–25*). So far, a number of encapsulin systems with different cargo enzymes have been studied and found to be involved in varied metabolic functions. Cargo enzymes include ferroxidases involved in iron storage (*7, 18, 19, 26–28*), peroxidases involved in oxidative stress resistance (*24, 29–31*), cysteine desulfurases involved in sulfur storage (*32, 33*), and terpene cyclases involved in terpenoid biosynthesis (*34, 35*). Enzyme encapsulation inside encapsulin shells can convey a number of distinct advantages, such as the formation of an optimized reaction environment, enzyme co-localization, intermediate sequestration, and the exclusion of unwanted proteins and molecules (*36*). Furthermore, the encapsulin shell may also serve to regulate cargo enzymes by modulating the flux of small molecule metabolites through its pores (*5, 21, 37*), whilst also stabilizing the oligomeric state of enzymatic cargo (*31*). Encapsulins are found in at least 35 prokaryotic phyla and have been classified into four distinct families based on operon structure and shell protein phylogeny (*6, 33*). Family 2, the most abundant class of encapsulins, is further subdivided into Family 2A and Family 2B based on the absence or presence, respectively, of a putative cyclic nucleotide binding domain (CBD) inserted into the E-loop of the HK97 fold encapsulin shell protein (*6*). Interestingly, only present within Family 2B are encapsulin operons encoding two discrete encapsulin shell proteins, with a diversity of associated cargo types (*5, 6*). There is no precedent for a multi-component encapsulin shell formed from more than one HK97 fold shell protein.

In this work, we report the characterization of a Family 2B two-component encapsulin shell identified in *Streptomyces lydicus* (Sl). Through heterologous expression and purification studies, we show the differential assembly behavior of the two shell components – Sl-Enc1 and Sl-Enc2 – and demonstrate their ability to co-assemble into mixed shells with variable shell composition. We determined the structures of both *S. lydicus* shell proteins using cryo-electron microscopy (cryo-EM). Using 3D classification analyses, we confirm co-assembly into mixed shells and highlight the irregular tiling and random incorporation of the two shell proteins into two-component shells. Our work expands the known assembly modes of HK97 fold proteins and lays the foundation for future studies aimed at elucidating the impact of two-component shells on encapsulin function.

## Results

### Two-component encapsulin operons are diverse and widespread in bacteria

To investigate the diversity and distribution of two-component encapsulin systems, we carried out a computational search to initially identify all Family 2B encapsulin shell genes encoded within 10 kb of each other in a given genome. Our search yielded 503 two-component operons representing 21% of all Family 2B systems (Fig. 1C). Five distinct operon types could be identified based on the identities of co-encoded putative cargo proteins (*5, 6*). The five operon types encode terpene cyclases (TCs), prenyltransferases (PTs), TCs and PTs together (TC/PT), xylulose kinases (XKs), and cysteine desulfurases (CDs), in decreasing order of prevalence. In addition, a number of two-component operons do not encode any obvious cargo (Other). The two encapsulin shell genes are generally clustered together with conserved gene synteny. We define the shell protein encoded by the first (upstream) gene as Enc1 and the shell protein encoded by the second (downstream) gene as Enc2. Most cargos are encoded directly up- or downstream of Enc1-Enc2, except for in TC/PT operons where putative TC cargos can be located between the Enc1 and Enc2 genes while putative PT cargos are found directly downstream of Enc2.

Two-component operons are distributed across five bacterial phyla with more than 75% of systems found in the phylum Actinobacteria which also exhibits the largest diversity of operon types (Fig. 1D). The other two phyla containing substantial numbers of two-component systems – Proteobacteria and Cyanobacteria – primarily encode TC-containing operons. Phylogenetic analysis of all encapsulin shell proteins identified in two-component operons highlights clustering within a phylum primarily based on cargo type (Fig. 1E). Enc1s within the same phylum sharing the same cargo type are more closely related to one another than to the Enc2s found in their respective operons. This observation may indicate a shared evolutionary trajectory of Enc1s, separate from Enc2s, and vice versa, prior to the formation of two-component operons. It may further suggest distinct and conserved roles for Enc1s and Enc2s in two-component systems.

Sequence analysis reveals two conserved diagnostic sequence differences – diagnostic motif 1 and 2 – between Enc1 and Enc2 shell proteins (Fig. 1F). These short sequence stretches represent the only conserved substantial differences between Enc1 and Enc2. Using sequence alignments, the sequence identities of the HK97 domains of all Enc1s, all Enc2s, and all two-component shell proteins combined (Enc1s and Enc2s) were determined to be 74.6%, 70.6%, and 70.3%, further confirming the phylogenetic clustering observed above (Fig. 1G). In addition, alignments focused only on the CBDs revealed a similar sequence identity trend – Enc1-CBDs: 51.9%, Enc2-CBDs: 49.0%, and Enc1/Enc2-CBDs: 48.5% – but with overall lower identities, highlighting that CBDs generally represent the most variable regions in Family 2B two-component shell proteins including Sl-Enc1 and Sl-Enc2 (fig. S1).

### Heterologous expression of individual shell components highlights differential assembly behavior

Here, we focus on the two encapsulin shell proteins found in a putative XK two-component system from *S. lydicus* – Sl-Enc1 and Sl-Enc2. We first sought to determine whether Sl-Enc1 and Sl-Enc2 would individually assemble into encapsulin shells. Separate heterologous expression of Sl-Enc1 and Sl-Enc2 in *E. coli* BL21 (DE3), followed by affinity purification and size exclusion chromatography, yielded soluble purified protein (Fig. 2A). Negative stain transmission electron microscopy (TEM) analysis of purified Sl-Enc1 and Sl-Enc2 highlighted that Sl-Enc1 did not assemble into regular encapsulin nanocompartments (Fig. 2B). In contrast, Sl-Enc2 formed characteristic ca. 28 nm shells, consistent with other T=1 Family 2B encapsulins (*35*). Native PAGE analysis further showed that Sl-Enc1 does not yield a single high molecular weight band – as observed for Sl-Enc2. Rather, a broad smear is observed, confirming that Sl-Enc1 does not assemble into homogeneous protein shells, forming irregular aggregates instead (Fig. 2C). This is surprising as Sl-Enc1 and Sl-Enc2 exhibit 79% overall sequence similarity and 89% sequence similarity of their shell-forming HK97-fold domains (fig. S1). The abovementioned diagnostic Enc1 and Enc2 sequence motifs, also found in Sl-Enc1 and Sl-Enc2, as well as the more variable CBDs may contribute to the observed differential assembly behavior of Sl-Enc1 and Sl-Enc2.

**Figure 2.**
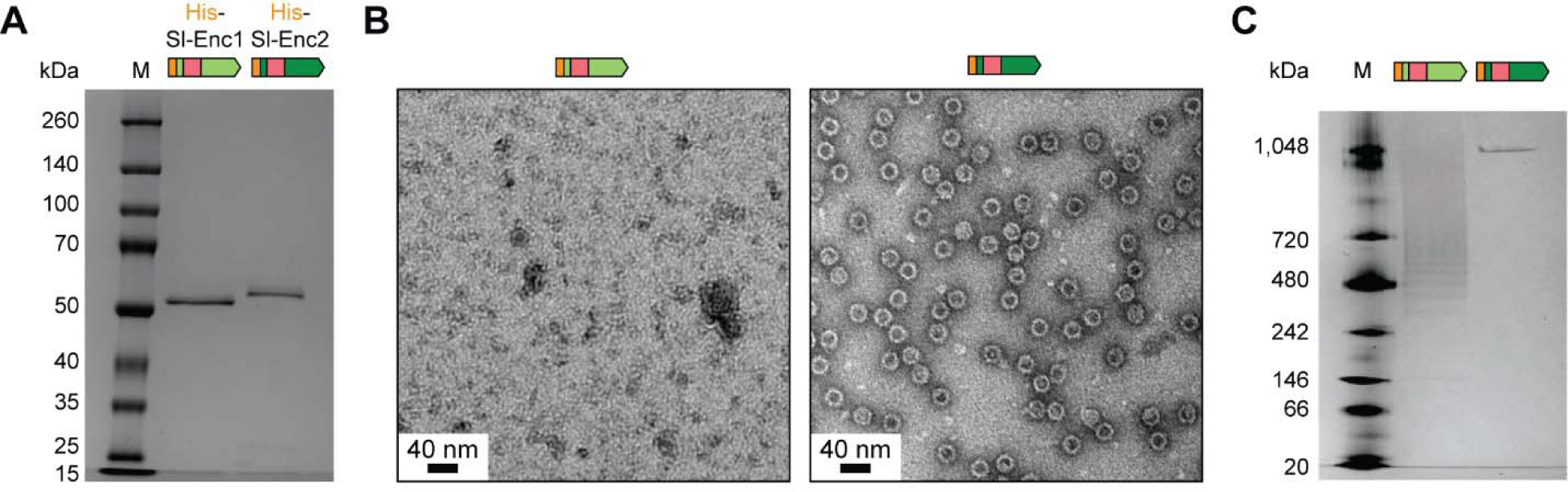
Separate analysis of Sl-Enc1 and Sl-Enc2. (**A**) SDS-PAGE analysis of individually expressed and purified Sl-Enc1 and Sl-Enc2. M: molecular weight marker. (**B**) Negative stain TEM micrographs of purified Sl-Enc1 and Sl-Enc2. (**C**) Native PAGE analysis of purified Sl-Enc1 and Sl-Enc2.

### Sl-Enc1 and Sl-Enc2 co-assemble into compositionally variable two-component shells

To determine if Sl-Enc1 and Sl-Enc2 can co-assemble to form mixed encapsulin shells, despite the fact that Sl-Enc1 cannot form shells on its own, we pursued a differential His-tagging strategy. We constructed an expression plasmid encoding an Sl-Enc1-Sl-Enc2 two-gene operon with Sl-Enc1 carrying an N-terminal His-tag. As Sl-Enc1 cannot assemble into shells by itself, this tagging strategy would allow us to utilize affinity purification and size exclusion chromatography to confirm shell formation, followed by protein identification to investigate the presence of both Sl-Enc1 and Sl-Enc2. Heterologous expression and purification of Sl-Enc1-Sl-Enc2 was carried out as described above. Clear encapsulin shells could be observed after purification via negative stain TEM (Fig. 3A) and native PAGE (Fig. 3B). The shells were of similar size and appearance compared to one-component Sl-Enc2 shells (Fig. 2B). SDS-PAGE analysis showed the presence of two distinct bands (Fig. 3C), identified as Sl-Enc1 (lower band) and Sl-Enc2 (upper band) via tryptic digest and LC-MS analysis. Repeat expressions and purifications of Sl-Enc1-Sl-Enc2 resulted in fairly consistent apparent ratios of Sl-Enc1 and Sl-Enc2 (ca. 60% Sl-Enc1 and 40% Sl-Enc2) (Fig. 3C). Because only Sl-Enc1 is His-tagged, Sl-Enc1 must be incorporated into a mixed shell since it cannot form shells by itself and because one-component untagged Sl-Enc2 shells – which are likely formed as well – would have been removed during affinity purification. Notably, the incorporation of Sl-Enc1 did not alter the size or appearance of mixed encapsulin shells compared with homogeneous Sl-Enc2 shells. As Sl-Enc1 and Sl-Enc2 are highly similar in sequence, Sl-Enc1 appears to be able to replace Sl-Enc2 subunits in mixed shells. Following a similar differential His-tagging strategy as described above, a two-gene operon with only Sl-Enc2 carrying an N-terminal His-tag was constructed, expressed, and purified to determine if Sl-Enc1 could be included in shells containing primarily Sl-Enc2. Shell assembly was confirmed using TEM and native PAGE (fig. S2, A and B). Although only a single major band was observed via SDS-PAGE analysis, LC-MS and tryptic digest of the excised gel band demonstrated that Sl-Enc1 was present (fig. S2C). This suggests that Sl-Enc1 is a minor component of this mixed shell sample, although, given that the relative protein ratios are estimated at a bulk level, it does not exclude the possibility of some individual shells having high incorporation of Sl-Enc1. Altogether, our differential His-tagging experiments establish that Sl-Enc1 incorporation into the encapsulin shell is variable and can range from 0% up to approximately 60% incorporation.

**Figure 3.**
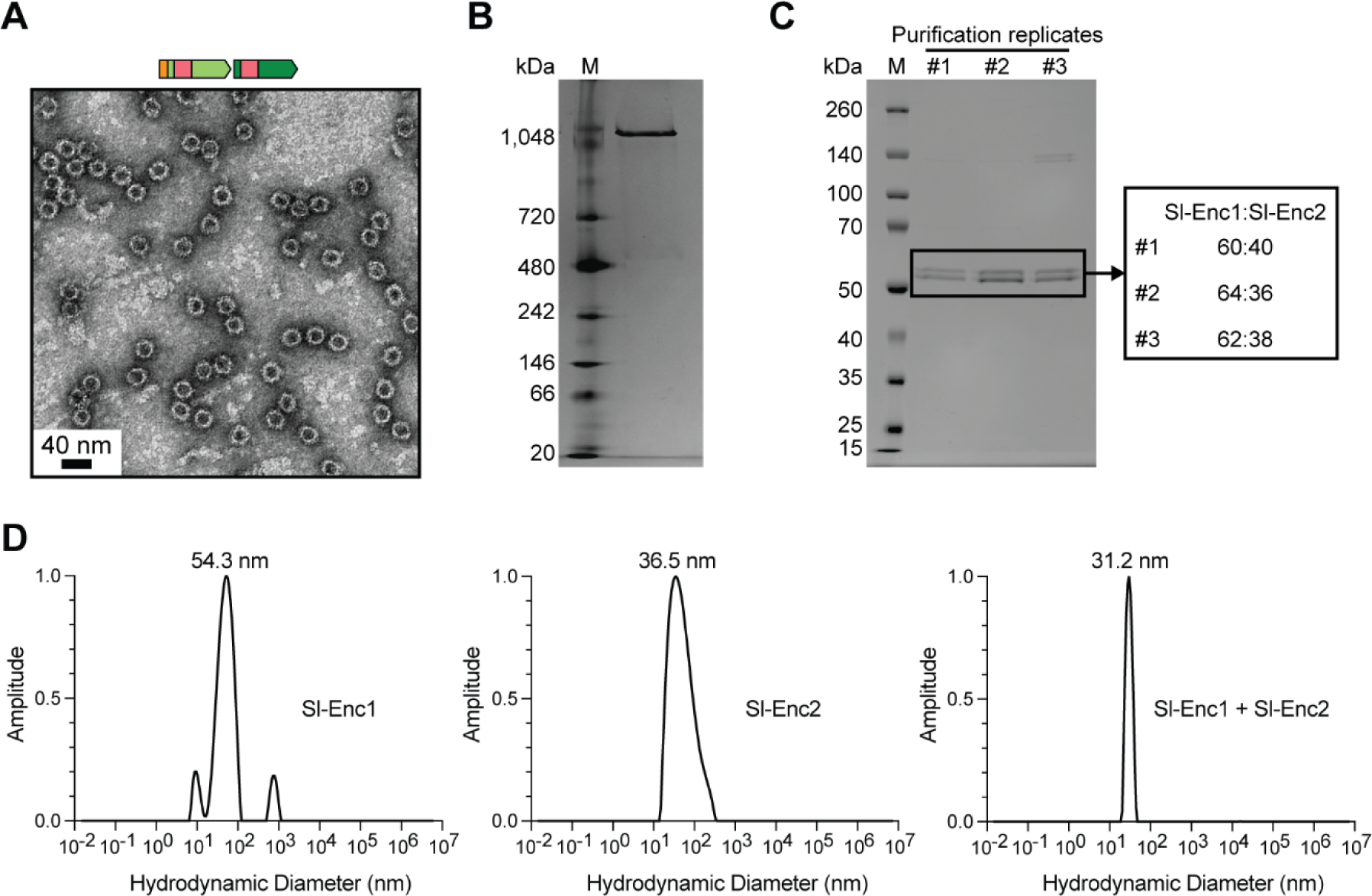
Analysis of mixed Sl-Enc1-Sl-Enc2 shells. (**A**) Negative stain TEM micrograph of purified Sl-Enc1 majority mixed shells. (**B**) Representative native PAGE analysis of an Sl-Enc1 majority mixed sample. (**C**) SDS-PAGE analysis of three replicate Sl-Enc1 majority mixed samples. The Sl-Enc1 to Sl-Enc2 ratio based on gel densitometry is shown on the right for individual purification replicates. (**D**) Dynamic light scattering (DLS) analysis of individually purified Sl-Enc1 (left) and Sl-Enc2 (middle) as well as of an Sl-Enc1 majority mixed sample (right). PDI: polydispersity index.

A potential preference for mixed shell formation is suggested by dynamic light scattering (DLS) analysis of purified samples with mixed shells exhibiting the narrowest size distribution and lowest polydispersity index (PDI) values (Fig. 3D). The broad distribution and high PDI of Sl-Enc1 further confirms its inability to form homogeneous assemblies. Thermal ramps in combination with DLS and static light scattering (SLS) showed that mixed samples possess the highest melting temperature (T_m_) of all analyzed samples, confirming that mixed shells are more stable than homogeneous Sl-Enc2 shells (fig. S3).

### Single particle cryo-EM analysis confirms mixed shell assembly and suggests irregular tiling of two-component shells

To gain a deeper understanding of the composition and structure of two-component shells, we performed single-particle cryo-EM analysis. In particular, we focused our efforts on two mixed samples with one of them exhibiting an excess of Sl-Enc1 (fig. S4A) and the other showing an excess of Sl-Enc2 (fig. S4B). Initially, we carried out consensus icosahedral (I) refinements and obtained symmetry-averaged cryo-EM maps at 2.58 Å (Sl-Enc1 majority) and 2.59 Å (Sl-Enc2 majority) (fig. S5 and S6). We found that protomer excess resulted in consensus maps exclusively representing the majority component. That is to say, the Sl-Enc1 majority sample yielded a symmetry-averaged Sl-Enc1 shell map while the Sl-Enc2 majority sample resulted in a symmetry-averaged Sl-Enc2 shell density. As Sl-Enc1 cannot form shells on its own, the obtained symmetry-averaged Sl-Enc1 map has to be treated as artifactual. However, we were able to use this map to build an atomic model of the Sl-Enc1 protomer (Fig. 4A). As mentioned above, Sl-Enc2 can assemble into complete shells by itself (Fig. 2B). Therefore, we used the symmetry-averaged Sl-Enc2 map to build a model for a complete homogeneous Sl-Enc2 shell. The Sl-Enc2 shell is 28 nm in diameter, consists of 60 identical protomers, and exhibits T=1 icosahedral symmetry, similar to recently described Family 2B encapsulins (Fig. 4B) (*35*). Both Sl-Enc1 and Sl-Enc2 protomers consist of two domains, a shell-forming canonical HK97-domain, conserved in all encapsulins, and a CBD domain inserted into the E-loop of the HK97-domain, only found in Family 2B encapsulins. The HK97-fold portions of both Sl-Enc1 and Sl-Enc2 exhibited high quality cryo-EM densities which allowed confident atomic model building. However, both CBDs exhibited weaker and lower resolutions densities, likely due to their inherent flexibility as highlighted by local resolution analysis (fig. S5 and S6). To create complete Sl-Enc1 and Sl-Enc2 protomer models, we utilized AlphaFold predictions of the two CBDs, which fit the obtained lower resolution cryo-EM CBD maps very well and combined them with our HK97-fold domain models. While the overall structures of the Sl-Enc1 and Sl-Enc2 protomers are very similar, two major differences could be observed. First, Sl-Enc1 exhibits a clearly defined 23-residue N-arm, not present in the Sl-Enc2 protomer. Similar N-arms have been described in viral HK97-fold capsid proteins in addition to Family 2A encapsulins, but not Family 2B systems (*8, 32, 33*). In viruses, N-arms usually mediate inter-protomer interactions, can often adopt multiple dynamic conformations, and increase the overall stability of the viral capsid (*2, 38–40*). The second major difference between the Sl-Enc1 and Sl-Enc2 protomers is that their CBDs show different relative orientations with respect to the HK97-domain with the Sl-Enc1 CBD being tilted by ca. 23° compared to the Sl-Enc2 CBD (Fig. 4A and fig. S7). Overall, the homogeneous Sl-Enc2 shell is similar to recently structurally characterized Family 2B encapsulins with externally displayed CBDs being positioned above all two-fold pores (*35*). The two-fold pores are large and elongated with dimensions of 14 by 60 Å and partially occluded by the CBDs (Fig. 4, C and D). The absence of any resolvable N-arms in the homogeneous Sl-Enc2 shell contributes to the large dimensions of these pores. Two additional types of pores are present in the Sl-Enc2 shell at the five- and three-fold axes of symmetry with diameters of 6 and 3 Å, respectively.

**Figure 4.**
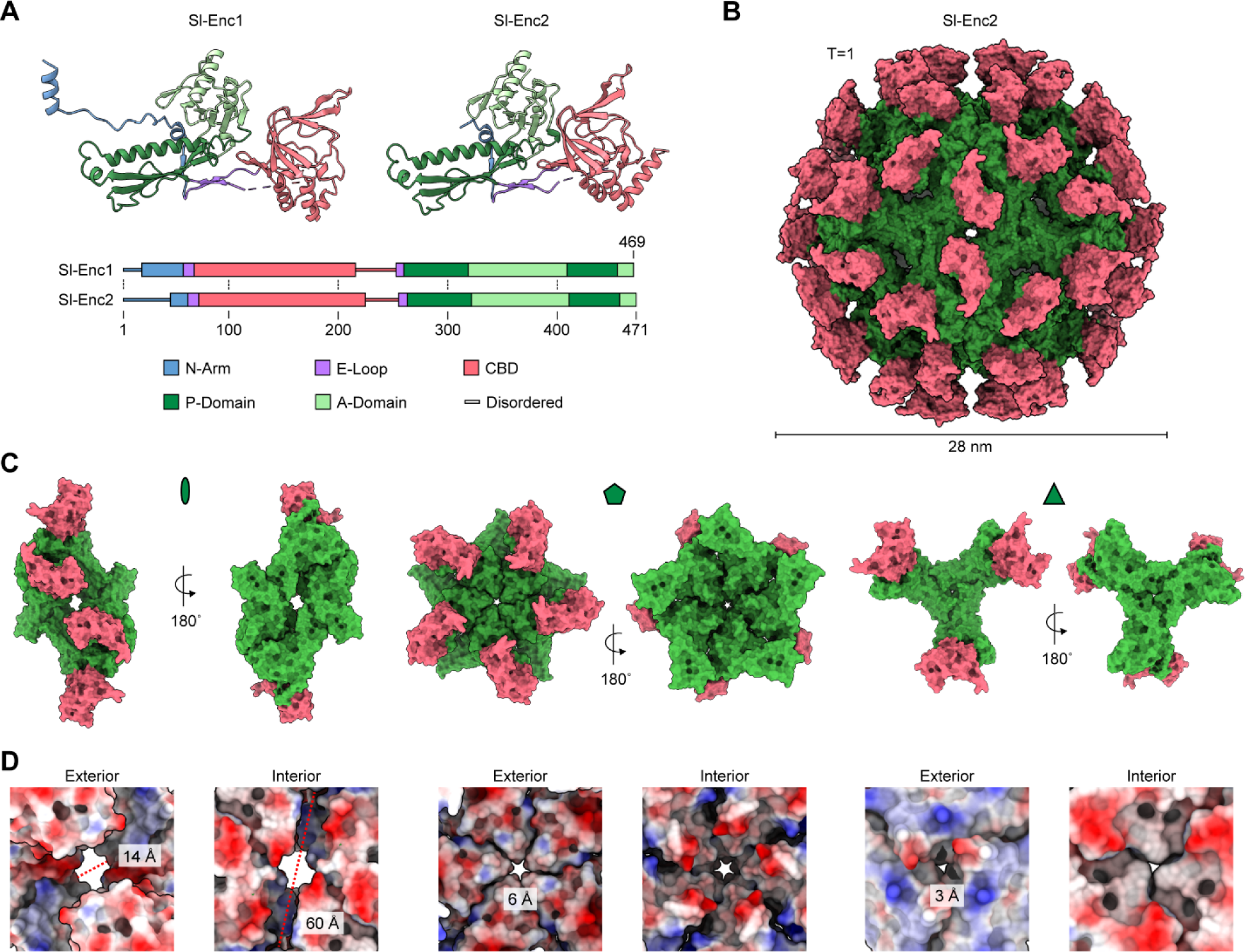
Structural analysis of mixed shells via single particle cryo-EM. (**A**) Protomer structures of Sl-Enc1 and Sl-Enc2. Structures colored by domain. (**B**) Homogeneous Sl-Enc2 shell with T=1 icosahedral symmetry. HK97-domain colored green. CBD colored pink. (**C**) Exterior and interior views of the two- (left), five- (middle), and three-fold (right) interactions found in the homogeneous Sl-Enc2 shell. (**D**) Two- (left), five- (middle), and three-fold (right) pores of the homogeneous Sl-Enc2 shell shown as electrostatic surface. Both the exterior and interior are shown with pore diameters highlighted.

C1 refinements of mixed samples generally yielded low resolution maps of poor quality suggesting substantial compositional heterogeneity. To visualize both protomer types – Sl-Enc1 and Sl-Enc2 – in mixed shells, we carried out masked 3D classifications followed by masked local refinements on symmetry-expanded particles (I symmetry) (fig. S8 to S11). We focused on the Sl-Enc1 majority sample where all shells can be assumed to be mixed as homogeneous Sl-Enc1 shells were not observed experimentally (Fig. 2B). We created focus masks for the four unique interactions each protomer within a Family 2B shell must engage in (Fig. 5A). It was our goal to identify all protomer interactions present in mixed two-component shells. We designed masks to completely encompass the HK97-domains of the two interacting protomers, omitting the CBDs due to their intrinsic conformational flexibility. We focused our analysis on two distinct interactions across the two-fold axis of symmetry (dubbed 2-fold (pore) and 2-fold (P-domain)), one across the three-fold axis, and one across the five-fold axis (Fig. 5B). In principle, four distinct interactions are possible for all investigated interfaces: Sl-Enc1/Sl-Enc1, Sl-Enc2/Sl-Enc2, Sl-Enc1/Sl-Enc2, and Sl-Enc2/Sl-Enc1, with the two mixed protomer arrangements representing the same but mirrored interaction. We were able to identify classes representing all four interactions for all masked 3D classifications (Fig. 5C and fig. S12 to S15). As the Sl-Enc1 and Sl-Enc2 protomers are highly similar in sequence and structure, we utilized one of the abovementioned diagnostic sequence motifs (diagnostic motif 1: Sl-Enc1: T_337_VWK_340_ vs Sl-Enc2: R_340_RRK_343_) to differentiate between Sl-Enc1 and Sl-Enc2 (Fig. 1F and Fig. 5D). Our results suggest that mixed two-component shells exhibit irregular tiling of protomers with all possible permutations allowed. We further found that one of the investigated two-fold interactions – 2-fold (pore) – exhibits additional heterogeneity with respect to the presence or absence of resolved N-arms (Fig. 6A). Specifically, two distinct symmetrical N-arm states could be observed in 3D classifications for Sl-Enc1/Sl-Enc1 interactions with N-arms either clearly resolved or completely absent. Symmetrical Sl-Enc2 or mixed interactions never exhibited resolved N-arms in our 3D classes. Thus, Sl-Enc1 protomers can exist in two different states, one with clearly resolved N-arms – as also observed in the symmetry-averaged consensus refinement – and one with completely unresolved N-arms. The discernable N-arm states result in substantial differences in the size and charge of the two-fold pores (Fig. 6, A and B).

**Figure 5.**
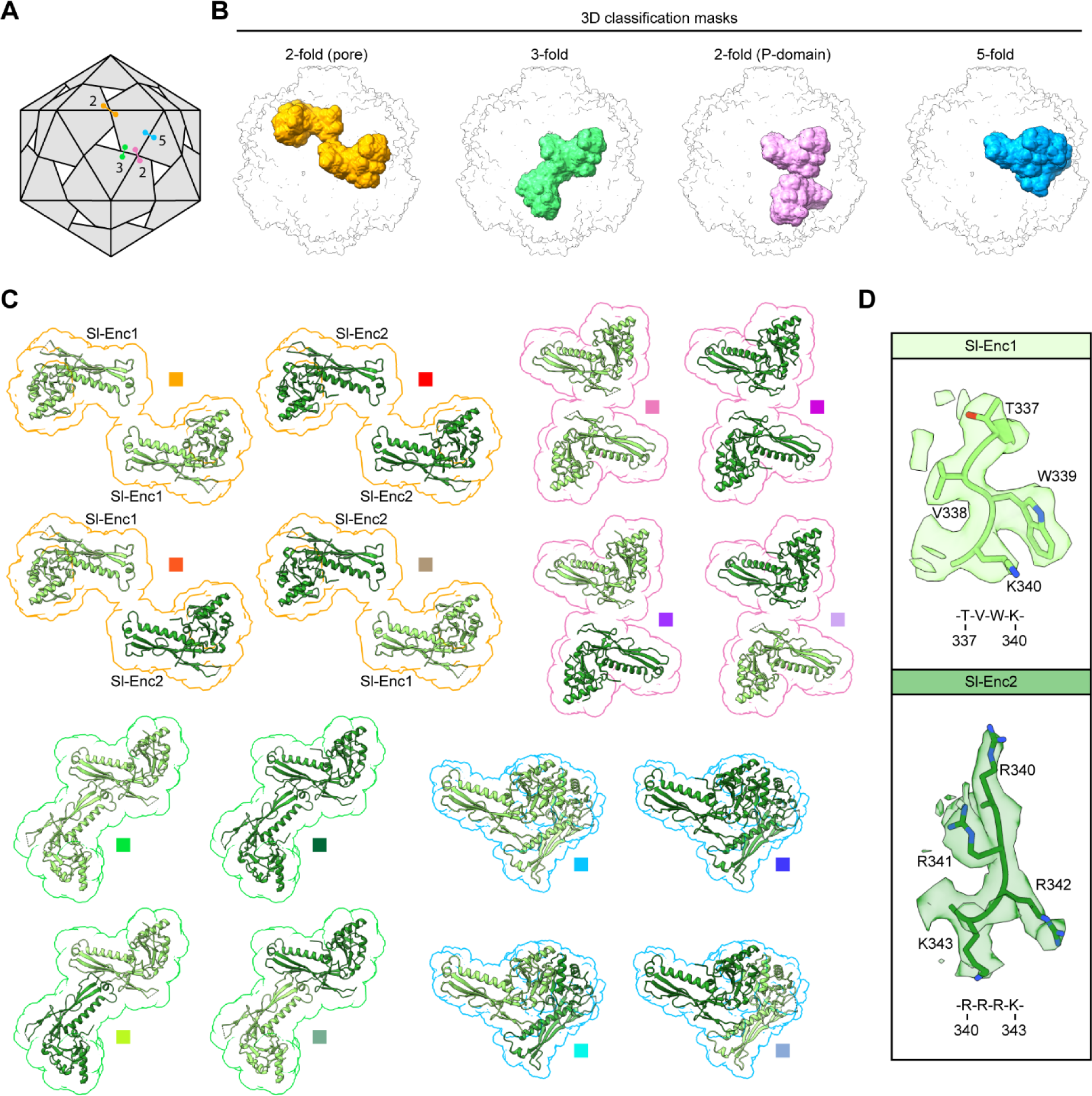
Protomer interaction analysis in mixed shells via 3D classification. (**A**) Schematic of a T=1 icosahedral protein shell highlighting the four unique interactions of a given protomer. Interactions are labelled by symmetry; orange: interaction across the 2-fold pore; green: interaction at the 3-fold pore; pink: 2-fold P-domain interaction; blue: 5-fold A-domain interaction. (**B**) 3D classification masks used to investigate all possible protomer interactions in mixed shells. (**C**) Observed protomer arrangements across the four unique interactions outlined in (A). Color scheme based on respective mask color. Sl-Enc1 shown in light green. Sl-Enc2 shown in dark green. (**D**) The diagnostic regions (diagnostic motif 1) within the Sl-Enc1 and Sl-Enc2 protomers used to unambiguously identify them. As an example, Sl-Enc1 from the Sl-Enc1-Sl-Enc1 2-fold P-domain interaction and Sl-Enc2 from the Sl-Enc2-Sl-Enc1 2-fold P-domain interaction are shown as cryo-EM density and atomic models with the respective sequence stretches shown below.

**Figure 6.**
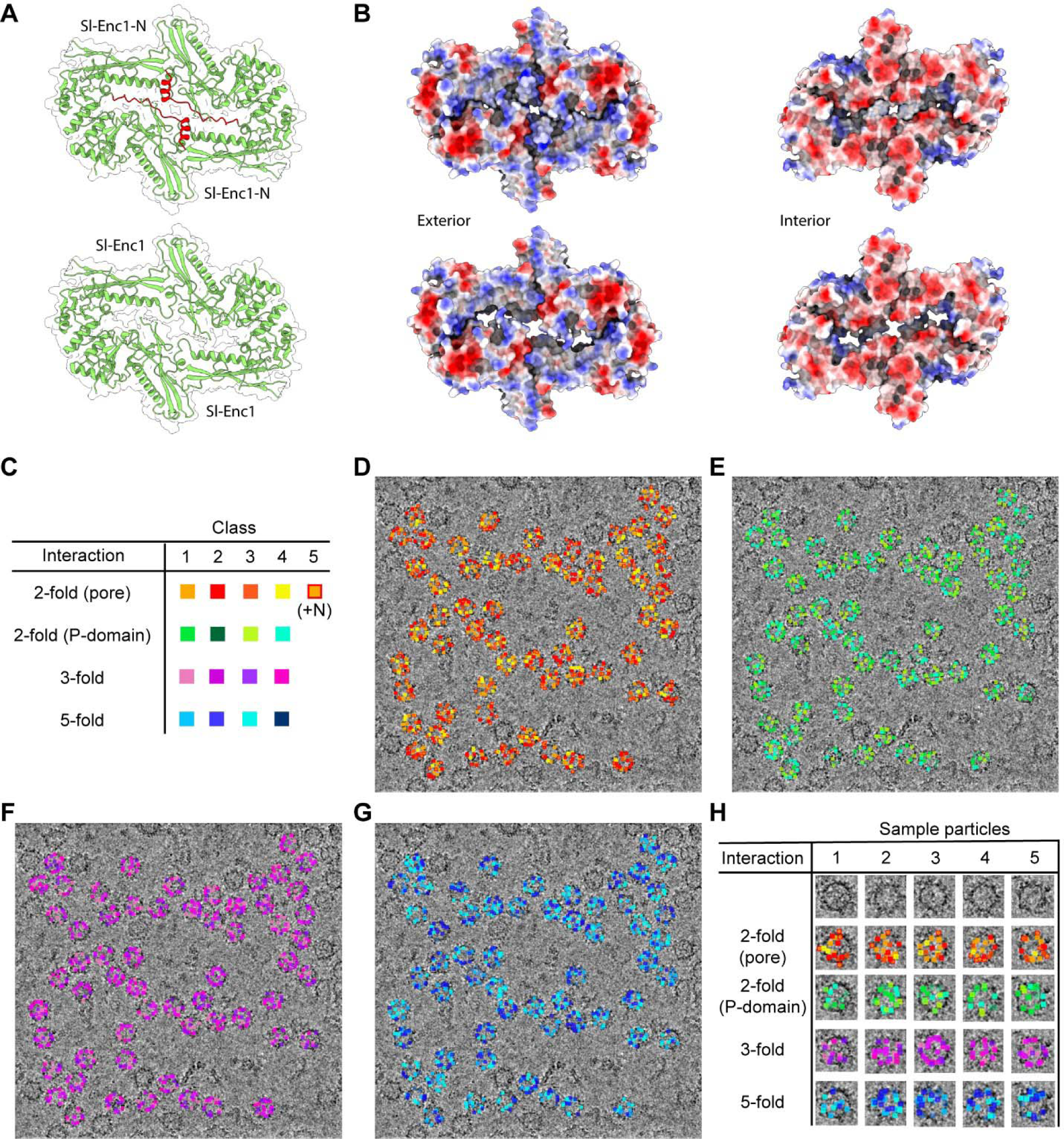
Two-fold pore variability and visualization of mixed shell heterogeneity. (**A**) Two-fold pore Sl-Enc1-Sl-Enc1 interactions observed via 3D classification analysis highlighting two resolved (top) or unresolved (bottom) N-arms (red). (**B**) Electrostatic surface representation of the two distinct two-fold pore Sl-Enc1-Sl-Enc1 arrangements. Top: two resolved N-arms; Bottom: unresolved N-arms. Left: exterior view. Right: interior view. (**C**) Color labelling scheme for the four unique interactions and protomer arrangements found across them used in subsequent panels. (**D**) Back-mapping of sub-particles contributing to the different classes identified for the 2-fold (pore) interaction via 3D classification onto a sample micrograph highlighting that all classes, representing unique protomer arrangements, can generally be found in each individual encapsulin particle. Each colored dot represents a single sub-particle sorted into one of the highlighted classes during 3D classification. (**E**) Back-mapping of sub-particles for the 2-fold (P-domain) interaction. (**F**) Back-mapping of sub-particles for the 3-fold interaction. (**G**) Back-mapping of sub-particles for the 5-fold interaction. (**H**) Five sample particles (1-5) from the sample micrograph shown above are highlighted with all the distinct protomer arrangements shown using the color scheme of panel (A). All identified protomer arrangements across the four unique interactions can generally be found within each individual encapsulin particle. As a subset of 10,000 particles was used for this analysis, not all interactions that exist within each particle will be present.

To show that all identified interactions and protomer arrangements are indeed present in most shells, we mapped a subset of 10,000 symmetry-expanded particles, contributing to the identified 3D classes, back onto the original micrographs (Fig. 6, C to G). We used a subset of particles for visual clarity. This analysis suggests that all 16 possible positional permutations across the four investigated interactions are likely present in all shells (Fig. 6H).

Taken together, our analysis highlights that two-component shells consisting of Sl-Enc1 and Sl-Enc2 are pseudo-icosahedral and highly irregular with all possible protomer arrangements allowed. This compositional heterogeneity results in variable pore properties for the two-, three-, and five-fold pores. Pore variability is further exacerbated by Sl-Enc1 being able to exist with or without resolved N-arms.

### Crosslinking analysis of two-component shells confirms heterogeneous two-fold interactions

To further investigate mixed shells, specifically with respect to the two-fold CBD interaction, analogous to the HK97-domain two-fold (P-domain) interaction discussed above, crosslinking studies followed by mass spectrometric analysis were carried out. Surface-exposed cysteines were introduced into the CBDs of either Sl-Enc1 (R195C) alone or both Sl-Enc1 and Sl-Enc2 (G198C) via site-directed mutagenesis (Fig. 7A). The two kinds of mutant mixed shells were still able to properly assemble as confirmed by negative stain TEM of purified samples (fig. S16, A to C). Site-selective crosslinking was carried out using a 17.8 Å cysteine-reactive double maleimide crosslinker (BM(PEG)_3_) (Fig. 7A). As both Sl-Enc1 and Sl-Enc2 natively contain one buried cysteine residue, control crosslinking experiments with native mixed shells were carried out. No crosslinking was observed in these control experiments with no higher molecular weight crosslinked bands visible on SDS-PAGE (Fig. 7B). For mutant mixed shells containing only the Sl-Enc1 cysteine (C195), a single crosslinked band could be observed on SDS-PAGE while two bands could be clearly visualized when both Sl-Enc1 (C195) and Sl-Enc2 (C198) cysteines were present (Fig. 7C). Subsequent tryptic digest and tandem mass spectrometric analysis of excised bands showed that the upper band, present in both samples, contains Sl-Enc1-Sl-Enc1 crosslinked peptides and the lower band, only present in the double cysteine (Sl-Enc1: C195 and Sl-Enc2: C198) mixed shells, contains both Sl-Enc1-Sl-Enc2 and Sl-Enc2-Sl-Enc2 crosslinked peptides (Fig. 7D, fig. S16D, and data S1). These results confirm our 3D classification results of the two-fold (P-domain) interaction and highlight that all possible interactions across this two-fold interface are present in mixed shells.

**Figure 7.**
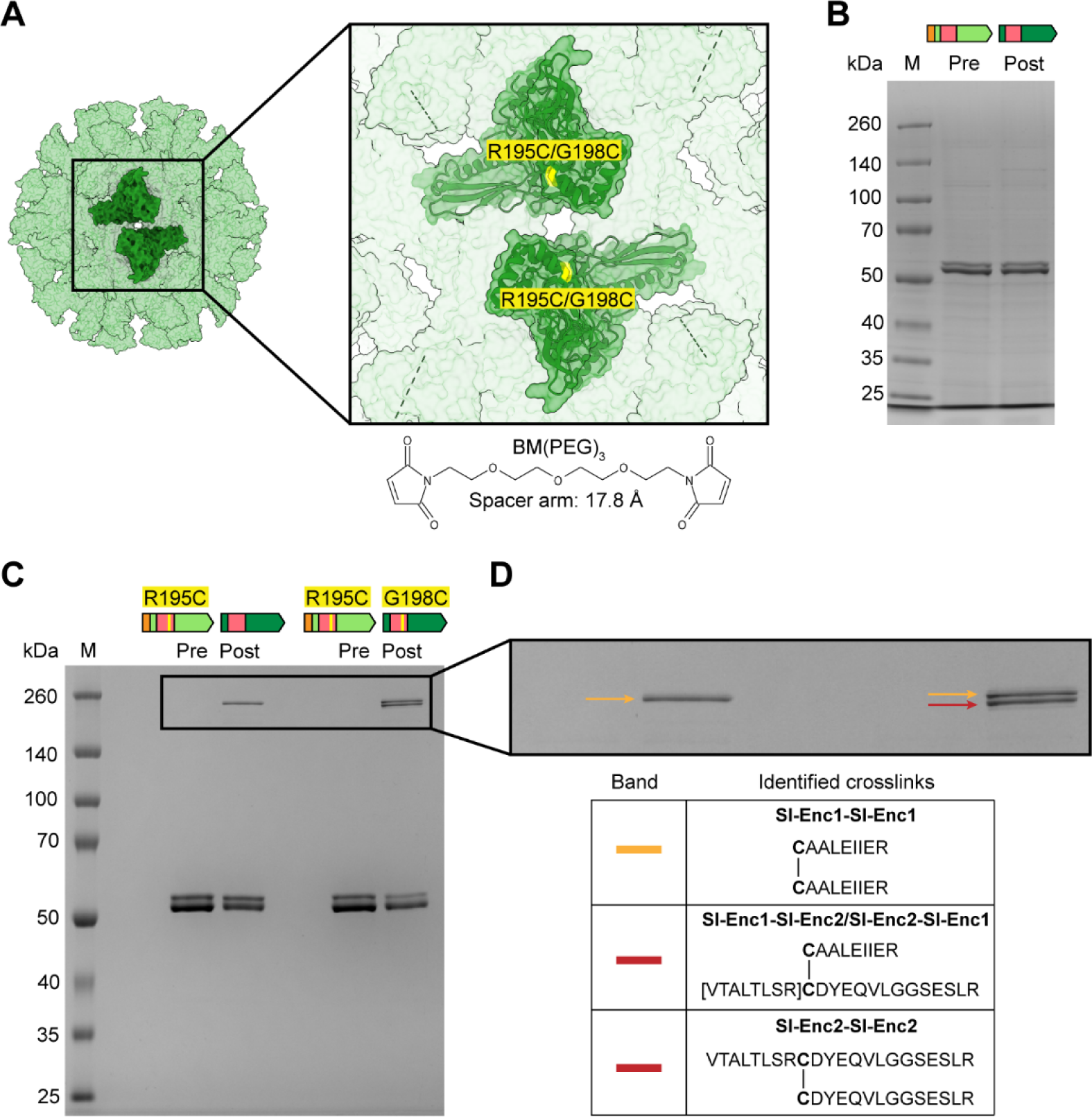
Site-specific crosslinking analysis of mixed shells. (**A**) Schematic of our crosslinking strategy based on CBD cysteine mutants (Sl-Enc1: R195C; Sl-Enc2: G198C). (**B**) SDS-PAGE analysis of a control crosslinking experiment using native mixed shells highlighting that no major higher molecular weight bands are present in the post crosslinking sample. (**C**) SDS-PAGE analysis of crosslinked mixed samples containing either one CBD cysteine (Sl-Enc1: R195C) or two CBD cysteines (Sl-Enc1: R195C and Sl-Enc2: G198C). The higher molecular weight bands observed in post crosslinking samples are outlined. (**D**) Mass spectrometric identification of crosslinks present in the higher molecular weight crosslinked bands. The identified crosslinks for each band are shown.

## Discussion

In this study, we report the structure and assembly principles of a two-component Family 2B encapsulin nanocompartment found in *Streptomyces lydicus*, highlighting its quasi-icosahedral geometry, variable shell composition, and irregular shell tiling. Our bioinformatic searches establish two-component encapsulin operons as widespread – 503 systems across five bacterial phyla, representing 21% of all Family 2B operons – and diverse – five distinct operon types with different cargo. Notably, the only operons to always encode two shell components are xylulose kinase (XK) systems. All other cargo types can be found in both two-component and single component operon arrangements (*6*). While this observation may be a result of the limited number and biased phylogenetic sampling of sequenced bacterial genomes, it could also indicate that some aspects of the two-component shell are essential for the proper functioning of specifically XK encapsulin systems. Our phylogenetic analysis further showed that Enc1s within a given phylum are more similar to each other than to Enc2s found within their respective operons (Fig. 1E). This could potentially hint at separate evolutionary histories of Enc1s and Enc2s which may have existed as single component encapsulin systems in the past. The strong gene synteny for Enc1 and Enc2 genes, observed in current two-component systems, would further suggest distinct and conserved roles for each shell component, supported by our experimental results demonstrating a preference for mixed shell formation (fig. S2 and S3).

We confirmed the co-assembly of Sl-Enc1 and Sl-Enc2 into mixed shells using a differential His-tagging strategy (Fig. 3) and further validated these findings with 3D classification analyses. We found that all possible protomer-protomer interactions are in principle allowed in mixed shells (Fig. 5, and Fig. 6, C to H). These findings demonstrate that both shell proteins can engage in all interactions across all interfaces which in turn implies that at the protomer-protomer interaction level, the sequence and structural differences between Sl-Enc1 and Sl-Enc2 do not restrict their interactions (Fig. 5). CBDs positioned across from each other arranged around the two-fold pore can similarly tolerate all four CBD pairings based on crosslinking studies (Fig. 7). These findings were surprising, as Sl-Enc1 is unable to form shells by itself (Fig. 2, B and C) implying that there exists some intrinsic limitation in its self-assembly capacity. This fact, in combination with Sl-Enc1’s propensity for aggregation rather than the formation of a single non-shell oligomeric state, may indicate that its inability to form homogeneous shells results from a cumulative structural effect. For example, slightly unfavorable angles between Sl-Enc1 protomer-protomer interfaces or the presence of too many defined N-arms – only observed for Sl-Enc1 – could potentially propagate if too many Sl-Enc1s are present in a given shell, resulting in an inability to form a closed compartment and a propensity towards aggregation. On the other hand, the observed preference for mixed shell formation may be partially attributed to the stabilizing effect of some, but not too many, defined Sl-Enc1 N-arms at the two-fold pore within mixed shells (Fig. 6). As shell protein features of Family 2B encapsulins are highly conserved, it seems reasonable to hypothesize that mixed shell formation represents a general feature of two-component encapsulin systems.

Using single particle cryo-EM, we found that the HK97-domain structures of Sl-Enc1 and Sl-Enc2 are nearly identical (Fig. 4A), with only three regions exhibiting substantial differences. One such difference is the presence of a resolved N-arm for Sl-Enc1 that was notably absent for Sl-Enc2. Similar N-arms, resulting in a chainmail-like topology, have also been observed in Family 2A encapsulins (*32, 33*) and many HK97 fold viral capsid structures and are believed to contribute to the generally high stability of HK97 fold assemblies. The other two regions of structural deviation between Sl-Enc1 and Sl-Enc2 – also detectable at the sequence level – were dubbed diagnostic sequence motifs (Fig. 1, F and G and fig. S17). One of these motifs – diagnostic motif 1 – is a stretch of four residues found in the A-domain located at the interface between two neighboring protomers within shell pentamers (Sl-Enc1: T_337_VWK_340_ vs Sl-Enc2: R_340_RRK_343_). Interestingly, these motifs are adjacent to the N-arms resolved for Sl-Enc1. The close proximity of this major difference between Sl-Enc1 and Sl-Enc2 to the N-arm could indicate that this diagnostic sequence motif may drive the preference for mixed shells by improving overall shell stability through incorporation of structured Sl-Enc1 N-arms into Sl-Enc2 shells. The other primary difference between the HK97-domains of Sl-Enc1 and Sl-Enc2 – diagnostic motif 2 – is found on the luminal side of the A-domain (diagnostic motif 2: Sl-Enc1: A_368_VDFHGHSLP_377_ vs Sl-Enc2: G_370_AELHGTDVR_379_). Since this motif is not located at a protomer-protomer interface, it is unlikely to impact shell assembly preferences. However, the positioning of this loop in the shell interior makes it a candidate for facilitating cargo loading (fig. S17). With two distinct shell components and two distinct internal loops, two different cargo-loading mechanisms could facilitate the loading of multiple cargo proteins with approximately constant cargo stoichiometries determined by shell composition.

One of the defining features of Family 2B encapsulins is the presence of an insertion domain of so far unknown function within the E-loop of the HK97 fold (*35*). It has been previously shown that even though annotated as a cyclic nucleotide-binding domain (CBD) – specifically a cyclic adenosine monophosphate (cAMP)-binding domain – exhibiting the expected CBD protein fold (PF00027), this insertion domain does not bind any cyclic nucleotides (*35*). This can be rationalized by some of the generally conserved CBD binding site residues being divergent in Family 2B CBDs. It was suggested that some other unidentified small molecule ligands may bind to encapsulin CBDs instead (*35*). The CBDs in Sl-Enc1 and Sl-Enc2 are the most variable regions of these shell proteins as highlighted by their low sequence identities (Fig. 1G). Further, the low local resolution observed via cryo-EM indicates a high degree of CBD conformational flexibility or mobility (fig. S5 and S6). The binding sites of the two CBDs in Sl-Enc1 and Sl-Enc2 also deviate from canonical CBD binding sites. Although the specific roles of CBDs remain unclear, their positioning above the major two-fold pore likely has implications for overall encapsulin function. It would be expected that the conformation and relative positioning of CBDs would greatly influence the level of occlusion at the two-fold pore, thereby regulating metabolite flux through the shell, potentially in a ligand-dependent fashion.

There are few known polyhedral protein compartments with shells or capsids that require multiple major, non-accessory or -dispensable, components and incorporate these components in a variable manner, as observed in the two-component encapsulin shell characterized in this work. One example are bacterial microcompartments (BMCs), a widespread class of bacterial protein compartments able to sequester multiple enzymes involved in carbon fixation or nutrient utilization (*41*). BMC shells are composed of three types of oligomers: hexamers (PF00936) that comprise most of the icosahedral faces (*42*), trimers (pseudo-hexamers) that influence the curvature and size of the compartment, also found within the icosahedral faces (*43, 44*), and pentamers (PF03319), necessary for forming closed shells, located at all shell vertices (*45*). Between three and eleven distinct shell components can be found across different predicted BMC operons (*46*). For one of the most well studied BMCs, the *Halothiobacillus neapolitanus* α-carboxysome, five distinct shell components are generally observed in natively isolated compartments (*47*). It has been suggested that this redundancy of shell components, especially for hexameric and trimeric oligomers, allows BMCs to adjust and fine-tune shell properties based on the dynamic needs of the host cell (*44*). For example, differences in pore charge and size likely influence the diffusion of small molecules across the BMC shell, and interactions with cargo proteins are usually mediated by specific shell proteins (*48, 49*). Since our studies of the *S. lydicus* two-component encapsulin were performed in the absence of cargo – no native cargo has yet been determined for XK Family 2B systems – it is possible that two-component encapsulins may benefit from the fine-tuning of shell properties for metabolite flux and having optimized interaction surfaces for cargo or external binders. Some hexameric BMC shell proteins have also been shown to form both homo- and hetero-hexameric oligomers of variable composition when heterologously expressed (*50, 51*). While this heterogeneity more closely mirrors the random tiling observed for the *S. lydicus* two-component shell, the functional role of BMC hetero-hexamers remains to be determined, especially since preliminary evidence is mixed regarding potential stability benefits (*50*).

Besides the encapsulin described in this work, bacteriophage T4 (T4) is another known example of an HK97 assembly to preferentially utilize two separate protomers (*52*). Structural data has demonstrated that one T4 capsid protein (gp24) forms homo-pentamers that cap the T4 vertices, while a second T4 capsid protein (gp23) forms homo-hexamers that co-assemble with pentamers into the T4 prolate head with icosahedral T=13 ends closing the central cylindrical section (*53, 54*). All other HK97 capsids or shells with a triangulation number greater than T=1 rely on the intrinsic flexibility of the HK97 fold protomer – especially the E-loop – to form both pentamers and hexamers from a single protomer. The two T4 capsid proteins (gp23 and gp24) have only 17% amino acid sequence identity (*16*), much lower than the two *S. lydicus* encapsulin shell proteins (fig. S1), although their structures are highly similar. Several A-domain mutations have also been identified that allow gp23 to form pentamers and substitute for gp24 (*54, 55*). No evidence has been found to suggest the presence of hetero-pentamers or - hexamers in the T4 capsid, in contrast to the *S. lydicus* two-component encapsulin which is composed of seemingly random hetero-pentamers with variable composition of the two shell components Sl-Enc1 and Sl-Enc2.

There are a number of potential functions provided by a two-component encapsulin shell that are difficult to realize using a single shell system. For example, the encapsulation of multiple cargos with distinct cargo encapsulation mechanisms can be easily envisioned in a two-component shell with distinct luminal shell binding sites for targeting peptides or cargo loading domains provided by two distinct shell components. Further, having more than one shell component may allow the fine-tuning of pore properties to optimize the metabolite flux of small molecules through the shell. While the function of CBDs found in Family 2B encapsulins remains to be determined, it has previously been hypothesized that small molecule ligand binding to externally displayed CBDs may modulate the properties of the shell or even the activity of encapsulated cargo in a dynamic fashion (*35*). Having two distinct CBDs present within a single shell may further expand this regulatory potential through the ability to bind two distinct ligands, thus integrating two separate regulatory signals.

In conclusion, we have demonstrated the existence of an HK97 fold protein compartment that preferentially uses two distinct shell components to form irregularly tiled encapsulin shells, expanding the known assembly modes of HK97 fold proteins. Our work lays the foundation for future studies aimed at elucidating the biological function of XK encapsulin systems and dissecting the functional logic of two-component encapsulins in general.

## Materials and Methods

### Bioinformatic and phylogenetic analysis

Family 2B encapsulin amino acid sequences were accessed from the UniProtKB database on March 30, 2023, by initiating a search for sequences containing the two Pfam families PF00027 and PF19307, resulting in 2,892 initial sequences. To identify genome neighborhoods for all initial hits, the Enzyme Function Initiative-Genome Neighborhood Tool (EFI-GNT) was used (*56, 57*). Two-component encapsulin systems were identified by filtering for Family 2B encapsulin genes with start codons within 10 kb of each other. Operons containing encapsulin sequences with less than 400 amino acids were categorized as fragments and removed from the dataset, resulting in a final dataset containing 1,006 UniProtKB accessions in 503 unique two-component Family 2B operons. Encapsulin sequences were then downloaded in FASTA format and aligned using MAFFT (mafft.crbc.jp/alignment) (*58*) with default settings. The alignment was forwarded to the Phylogenetic Tree function on the MAFFT server to generate a tree with the following settings: method-NJ with conserved sites (180 AAs), substitution model-JTT, heterogeneity among sites-ignore (alpha = infinity), bootstrap-1000. The phylogenetic tree was then visualized and annotated using iTOL V6.7.3 (*59*).

### Molecular biology and cloning

The *S. lydicus* encapsulin shell genes were codon-optimized for *E. coli* using the Integrated DNA Technologies (IDT) codon optimization tool and ordered as gBlock gene fragments from IDT (table S1). pETDuet-1 vector was linearized by digestion with PacI and NdeI and gene fragments were inserted into multiple cloning site 2 (MCS2) of the linearized plasmid via Gibson assembly using the New England BioLabs (NEB) Gibson Assembly Master Mix and following the manufacturer’s standard protocol. All gene modifications, including N-terminal His- and Strep-affinity tags, as well as cysteine modifications, were introduced to the *S. lydicus* encapsulin genes via primer extension with primers from IDT using an inverse PCR strategy.

Outward-pointing PCR primer pairs were designed to amplify the entire template with the desired sequence insertions or mutations (table s2 and s3). Linear plasmid fragments were phosphorylated with T4 polynucleotide kinase and ligated with T4 DNA ligase following the manufacturer’s protocol. The double cysteine mutant plasmid was constructed using Gibson Assembly to ligate two PCR-amplified fragments with the desired mutations. Invitrogen One Shot BL21(DE3) chemically competent *E. coli* cells were subsequently transformed with the respective assembled plasmids and validated via Sanger sequencing (Eurofins) (table s4).

### Protein expression and purification

Protein expression for all constructs was carried out using Terrific Broth (TB) media supplemented with 100 µg/mL ampicillin. Cell cultures were inoculated from overnight starter cultures at a ratio of 1:100 and grown at 37°C, 200 rpm until reaching an OD_600_ of 0.4-0.8. Then, protein expression was induced with 0.5 mM isopropyl β-D-1-thiogalactopyranoside (IPTG) and cells grown further at 18°C for 20-24 h. Cells were harvested by centrifugation at 8,000 rcf for 15 min at 4°C and cell pellets were either used immediately or stored at −80°C.

Cell pellets were thawed and resuspended in 3 mL of Tris Buffered Saline (TBS) (20 mM Tris, 300 mM NaCl, pH 8.0) per 1 g of wet cell weight. Lysozyme (0.5 mg/mL), DNase (1 µg/mL), Benzonase nuclease (25 units/mL), and SIGMAFAST EDTA-free protease inhibitor cocktail (1 tablet per 100 mL) were added to the resuspension buffer. Cell suspensions were incubated at 4°C while stirring for one hour. Cells were lysed by three consecutive passages through an EmulsiFlex-C3 homogenizer at 20,000 psi and subsequently centrifuged at 8,000 rcf for 15 min at 4°C. The soluble fraction was then applied to a 5 mL packed Ni-NTA resin bed (GoldBio) and equilibrated with 10 column volumes of TBS. Resin was washed with a stepwise gradient in increments of 30 mM imidazole up to 90 mM. TBS buffer containing 250 mM imidazole was used to elute the remaining protein bound to the Ni-NTA resin. Protein samples were analyzed for purity using SDS-PAGE with 14% Novex Tris-Glycine Gels (Thermo-Fisher Scientific). Fractions containing pure protein were pooled and subsequently concentrated and buffer exchanged using 50 kDa MWCO Amicon Ultra-15 mL centrifugal units with TBS. The concentrated protein sample was either stored at 4°C or immediately purified further via size exclusion chromatography (SEC). For SEC purification, the concentrated protein sample was centrifuged for 10 min at 10,000 rcf and then loaded onto an ÄKTA Pure FPLC system and purified using a Superose 6 10/300 GL column pre-equilibrated with TBS. For cysteine mutants, proteins were purified in the same way, but the buffer contained 2-5 mM tris(2-carboxyethyl)phosphine (TCEP) throughout all purification steps.

### Negative stain transmission electron microscopy (TEM)

Purified encapsulin samples were imaged via negative stain transmission electron microscopy (TEM) with a 100 kV Morgagni transmission electron microscope at the University of Michigan Life Science Institute. 200 Mesh Gold Grids coated with formvar-carbon film (FCF-200-Au, EMS) were glow-discharged using a PELCO easiGlow at 5 mA for 60 s to increase their hydrophilicy. To the glow-discharged grids, 3.5 µL of a 0.1 mg/mL protein sample in 20 mM Tris, 150 mM NaCl was applied and allowed to adsorb for 1 min before excess liquid was blotted away with Whatman filter paper. The grid was first washed with distilled water and then with 0.75% (w/v) uranyl formate before being floated on an 11 µL drop of 0.75% (w/v) uranyl formate for one min. After removing excess stain via blotting, samples were immediately imaged or stored at room temperature for later use.

### Dynamic light scattering (DLS) and thermal ramp experiments

DLS and thermal stability measurements were carried out using an Uncle instrument (Unchained Labs) at 25°C in triplicate. Purified protein samples were adjusted to a concentration of 0.02 to 0.2 mg/mL, centrifuged at 20,000 rcf for 10 min at 4°C, and then immediately analyzed via DLS. Melting temperature (T_m_) and aggregation temperature (T_agg_) analysis was conducted over a 25°C to 95°C thermal ramp at 1°C per min, measuring static light scattering (SLS) at 266 nm.

### Cryo-electron microscopy (cryo-EM)

#### Sample preparation

Purified Sl-Enc1 or Sl-Enc2 majority samples were concentrated to 2 mg/mL in 150 mM NaCl, 25 mM Tris pH 8.0. 3.5 µL of protein sample were applied to freshly glow discharged Quantifoil R1.2/1.3 Cu 200 mesh grids (EMS, Q225-CR1.3) and were frozen by plunging into liquid ethane using an FEI Vitrobot Mark IV (100% humidity, 22°C, blot force 20, blot time 4 s, wait time 0 s). The grids were immediately clipped and stored in liquid nitrogen until data collection.

#### Data collection

Cryo-EM movies for Sl-Enc1-His-Sl-Enc2 (Sl-Enc1 majority) were collected using a ThermoFisher Scientific Talos Arctica cryo-electron microscope operating at 200 kV equipped with a Gatan K2 direct electron detector. 1,658 movies were collected from a single grid using the Leginon (*60*) software package at a magnification of 45,000x, pixel size of 0.91 Å, defocus range of −0.8 µm to −1.8 µm, exposure time of 5 s, frame time of 200 ms, and a total dose of 45.5 e^-^/Å^2^. Cryo-EM movies of Sl-Enc1-His-Sl-Enc2-His (Sl-Enc2 majority) were collected using a ThermoFisher Scientific Krios G4i cryo-electron microscope operating at 300 kV equipped with a Gatan K3 direct electron detector with a BioQuantum energy filter operating at 20 eV slit width. 2,865 movies were collected from a single grid using the Leginon (*60*) software package at a magnification of 105,000x, pixel size of 0.84 Å, defocus range of −0.5 µm to −1.0 µm, exposure time of 3 s, frame time of 50 ms, and total dose of 54 e^-^/Å^2^.

#### Image processing

CryoSPARC v4.2.1 was initially used to process the Sl-Enc1 majority dataset (*61*). 1,658 movies were imported, motion corrected using patch motion correction, and the CTF-fit was estimated using patch CTF estimation. Exposures with CTF fits worse than 5 Å were discarded from the dataset, resulting in 1,541 remaining movies. 222 particles were manually picked to create templates for template-based particle picking. Template picker was used to identify particles and 77,425 particles were extracted with a box size of 440 pixels. Two rounds of 2D classification yielded 72,804 particles. Ab-initio reconstruction with 3 classes and I symmetry was then carried out. 72,760 particles were contained in the major class and were then used for heterogeneous refinement with 3 classes and I symmetry imposed. This yielded a major class with 69,590 particles which were then subjected to homogeneous refinement against the heterogeneous refinement map with I symmetry imposed, per-particle defocus optimization, per-group CTF parameterization, and Ewald sphere correction enabled, resulting in a 2.58 Å resolution consensus map. Local resolution was calculated in cryoSPARC.

For all subsequent analyses of the Sl-Enc1 sample, cryoSPARC v4.4.0+231114 was used. To investigate the compositional heterogeneity and allowed protomer arrangements within the Sl-Enc1 majority sample, the 69,590 particles used to create the consensus refinement were initially symmetry expanded (I symmetry), yielding 4,175,400 particles, of which 3,675,400 particles were – due to computational limitations – used for further downstream processing. Four separated masked 3D classification runs were carried out using 3,675,400 symmetry expanded particles with the following non-standard parameters: 16 classes, target resolution: 3.5, O-EM learning rate init: 0.75, O-EM learning rate halflife: 100, force hard classification, class similarity: 0.25. Four custom 3D classification masks were created for the four unique protomer interactions possible within a T=1 icosahedral shell using ChimeraX v1.6.1 and the volume tools job (dilation radius: 1, soft padding: 3) in cryoSPARC. All masks included only the HK97-domains, and not the CBD domains, of the two protomers in question. After manual inspection, select 3D classes were refined via masked C1 local refinements using the same masks as for the 3D classification jobs. The following non-standard parameters were used: use pose/shift gaussian prior, standard deviation of prior over rotation: 7, standard deviation of prior over shifts: 4, maximum alignment resolution: 0.05, re-center rotations each iteration, re-center shifts each iteration, number of extra final passes: 2, initial lowpass resolution: 6, force re-do GS split. To map symmetry-expanded particles of select 3D classes onto micrographs, the following protocol was used: first, a subset of 10,000 particles – created via the particle sets tool – from each 3D class of interest was re-centered using the volume alignments job in cryoSPARC. As the new center coordinates, the centers of the respective 3D classification masks were used which were measured in ChimeraX. The resulting re-centered particles were then re-extracted with a box size of 32 pixels. As these particles were exclusively used for visualization purposes, a small box size was sufficient. The re-extracted re-centered particles were then mapped onto micrographs and visualized via the inspect picks job. To create images of symmetry-expanded particles from multiple classes mapped onto the same micrograph, individually mapped micrographs were overlayed to create composite images.

CryoSPARC v4.2.1 was used to process the Sl-Enc2 majority dataset. 2,865 movies were imported, motion corrected using patch motion correction, and the CTF-fit was estimated using patch CTF estimation. Exposures with CTF fits worse than 5 Å were discarded from the dataset, resulting in 2,176 remaining movies. After manual picking and template creation, template picker was used to pick 134,284 particles which were subsequently extracted with a box size of 480 pixels. Two rounds of 2D classification yielded 116,239 particles. These particles were then used for heterogeneous refinement with 3 classes and I symmetry applied. 64,557 particles were contained in the major class and were then used for homogeneous refinement with I symmetry, per-particle defocus, and per-group CTF parameterization enabled. This yielded a 2.59 Å resolution consensus map. Local resolution was calculated in cryoSPARC.

#### Model building

Cryo-EM maps were initially opened in UCSF Chimera v 1.15 (*62*) and for the Sl-Enc2 majority dataset, an initial model of the *Streptomyces griseus* Family 2B encapsulin (PDB ID: 9BHU) was manually placed into the density and the fit further optimized using the Dock to Volume command. For the Sl-Enc1 majority dataset, an initial model of Sl-Enc2 was placed using the same procedure in USCF Chimera v 1.15. The resulting models were opened in Coot v 0.8.9.1 (*63, 64*) and manually refined against the maps using rigid body refinement, chain refinement, and iterative manual real space refinement along the length of the chain while also mutating chains to the respective protein (Sl-Enc1 or Sl-Enc2). The protomer models were then refined in Phenix v 1.20.1-4487 (*65–67*) using phenix.real_space_refine with default parameters. NCS was identified from the maps using phenix.map_symmetry, and NCS operators were applied using phenix.apply_ncs to make icosahedral models containing 60 copies of the respective protomer. These models were then refined using phenix.real_space_refine with NCS constraints, global minimization, and ADP refinements. The models were validated using the comprehensive validation tool via phenix.validation_cryoem. The models were further inspected and manually refined until satisfactory geometry and density fit statistics were reached.

### Protein crosslinking assays

Purified encapsulin samples containing cysteine point mutations at 0.1 mg/mL (1 mL) were buffer exchanged using a 100 kDa MWCO Amicon Ultra-15 mL centrifugal unit into 20 mM Tris, 150 mM NaCl, 5 mM EDTA at pH 7.3. A fresh 5 mM stock of 1,11-bis(maleimido)triethylene glycol crosslinker (BM(PEG)_3_, Thermo-Fisher Scientific) was then prepared by dissolving the reagent in dimethylformamide (DMF). The crosslinker solution was added to the protein sample at a five-fold molar excess and the reaction mixture incubated at room temperature for 1 h in the dark. After incubation, the reaction was quenched by adding dithiothreitol (DTT) to a 10 mM final concentration, followed by incubation for 15 min at room temperature. Crosslinked protein samples were then analyzed via SDS-PAGE with 10% Novex Tris-Glycine gels (Thermo-Fisher Scientific).

### Protein identification via mass spectrometry

Purified Sl-Enc1/His-Sl-Enc2 protein samples were analyzed via SDS-PAGE with a 14% Novex Tris-Glycine Gels (Thermo-Fisher Scientific). The SDS-PAGE gel was stained with Coomassie and the overexpressed bands at the correct molecular weights were excised and stored in water. The SDS-PAGE gel bands were transferred to the University of Michigan Biomedical Research Core Facilities (BRCF) Proteomics & Peptide Synthesis Core, where the protein was extracted from the SDS-PAGE gel and subjected to tryptic digest at 37°C for 4 h, quenched, and analyzed. The digested samples were then analyzed using a nano LC/MS/MS Waters HPLC system coupled to a Thermo-Fisher Scientific LTQ Orbitrap Velos Pro following standard MS/MS protocol. Data were searched using a local copy of Mascot (Matrix Science, UK).

### Mass spectrometric analysis

Crosslinked protein bands were excised from SDS-PAGE gels and transferred to the University of Michigan Department of Chemistry Mass Spectrometry facility for processing and analysis. Gel bands were washed with water and 80% acetonitrile to elute protein from the gel matrix. Protein was treated with dithiothreitol (DTT) and 2-iodoacetamide (IAA) to reduce the disulfide bonds and alkylate the protein. The sample was subsequently digested with trypsin for 4 h at 45°C. Samples were analyzed on a Thermo-Fisher Scientific Orbitrap Fusion Lumos Tribrid mass spectrometer. Peptides were separated with a C18 desalting column (Thermo Acclaim PepMap, 75 µm × 25 cm) and reconstituted in 0.1% formic acid. A 60 min gradient from 4% to 76% acetonitrile with 0.1% formic acid, flow rate = 300 nL/min was used. HCD-MSMS was applied in data-dependent mode for a 3 second cycle time for all ions above 5e4 with 2+ to 7+ charge; applied 34% HCD energy. Proteome Discover 2.2 software was used to process the data and identify crosslinked peptides. The *S. lydicus* peptide sequences with affinity tag and cysteine modifications were used as sequence inputs.

## Acknowledgments

We acknowledge the staff of the University of Michigan Department of Chemistry Mass Spectrometry facility, as well as the Proteomics & Peptide Synthesis Core, for technical assistance.

## Author contributions

Conceptualization: CAD, TWG; Methodology: CAD, MPA, TWG; Investigation: CAD, MPA; Visualization: CAD, MPA, TWG; Supervision: TWG; Writing—original draft: CAD; Writing— review & editing: CAD, TWG.

## Data and materials availability

Cryo-EM maps and atomic models for Sl-Enc1 and Sl-Enc2 have been deposited in the Electron Microscopy Data Bank (EMDB) and the Protein Data Bank (PDB) and are publicly available (EMD-44632, EMD-44603; PDB ID: 9BJE, PDB ID: 9BIX). All data are available in the main text or the supplementary materials.

