## Supplementary Material for "A two-component quasi-icosahedral protein nanocompartment with variable shell composition and irregular tiling"

**This PDF file includes:**

Figs. S1 to S17
Tables S1 to S5

**Other Supplementary Materials for this manuscript include the following:**

Data S1

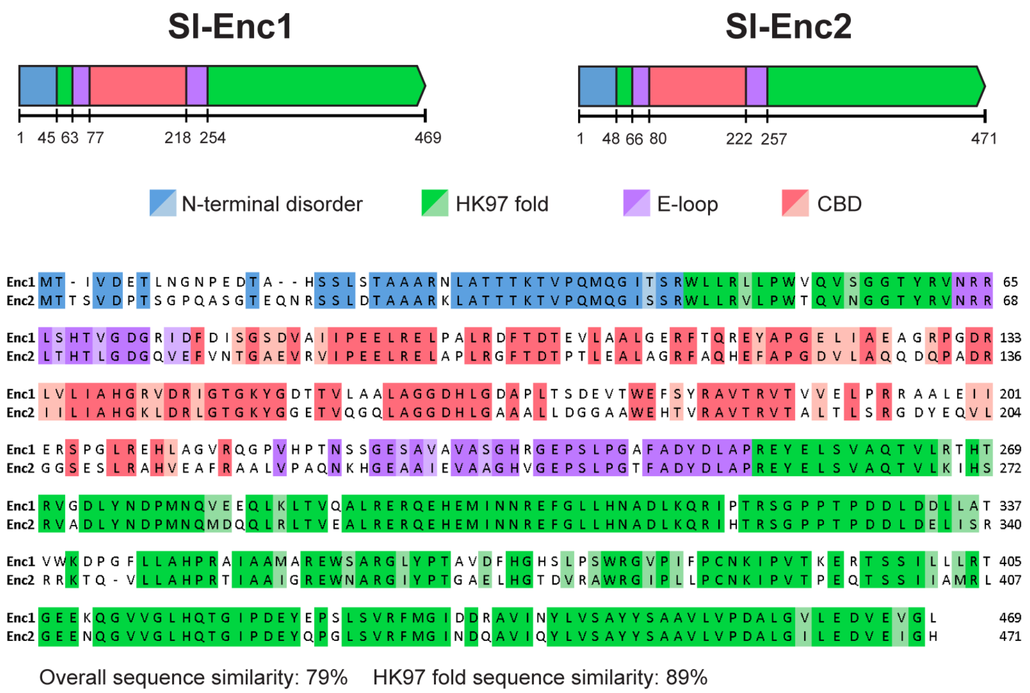


**Fig. S1. Sequence alignment of Sl-Enc1 and Sl-Enc2.** A sequence alignment of Sl-Enc1 and Sl-Enc2 generated via Clustal Omega is shown. Sequences are colored by domain.

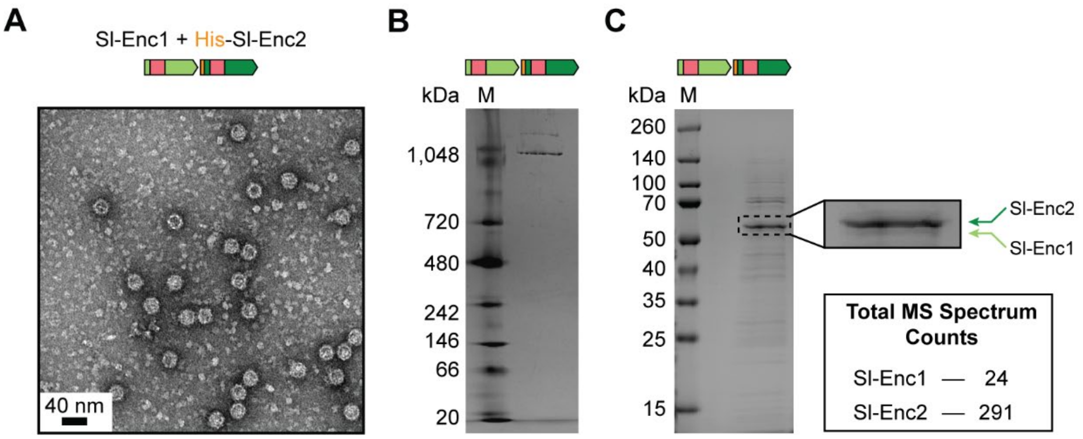


**Fig. S2. Analysis of Sl-Enc2 majority shells.** (**A**) Negative stain TEM micrograph of an Sl-Enc1 + His-Sl-Enc2 mixed shell sample purified via affinity purification and size exclusion chromatography. (**B**) Native PAGE analysis of an Sl-Enc1 majority mixed sample. M: molecular weight marker. (**C**) SDS-PAGE of purified Sl-Enc1 + His-Sl-Enc2. Outlined region on SDS-PAGE gel was excised and prepared for mass spectrometric analysis for protein identification. Total spectrum counts of Sl-Enc1 and Sl-Enc2 from mass spectrometry analysis are shown.

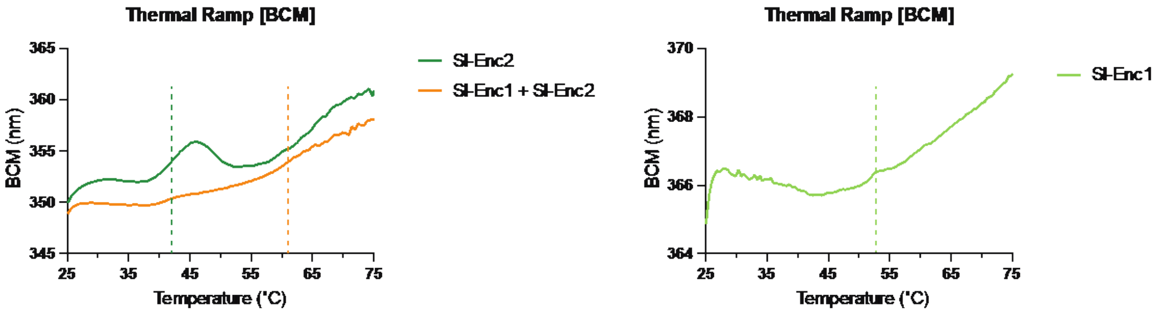


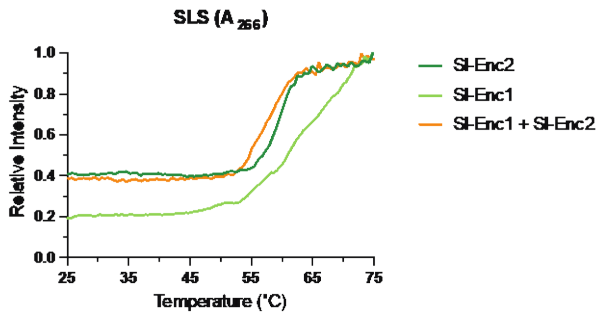


| | Average Tm | Average Tagg |
|---|---|---|
| Sl-Enc1 | 52.7 ± 5.8 °C | 51.5 ± 3.4 °C |
| Sl-Enc2 | 42.1 ± 2.0 °C | 53.7 ± 2.3 °C |
| Sl-Enc1 + Sl-Enc2 | 61.0 ± 2.2 °C | 51.5 ± 1.3 °C |

**Fig. S3. Thermal ramp data for Sl-Enc1 and Sl-Enc2.** (**A**) Thermal ramp data for Sl-Enc1, Sl-Enc2, Sl-Enc1 + Sl-Enc2. (**B**) SLS results for SL-Enc1, Sl-Enc1, and Sl-Enc1 + Sl-Enc2. (**C**) Average $T_m$ and $T_{agg}$ for Sl-Enc1, Sl-Enc2, and Sl-Enc1 + Sl-Enc2.

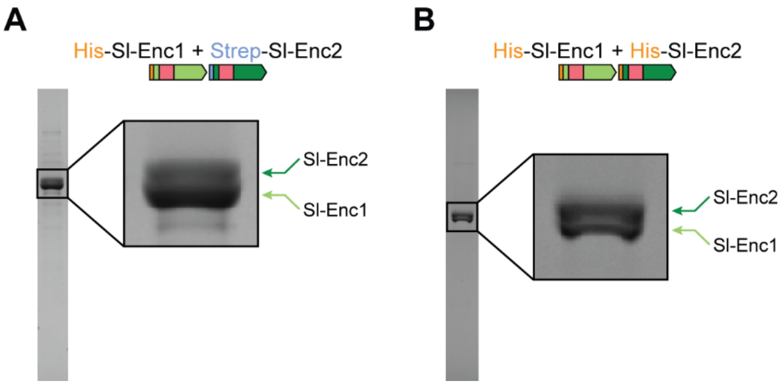


**Fig. S4. SDS-PAGE analysis of Sl-Enc1 majority and Sl-Enc2 majority samples subjected to cryo-EM analysis.** (**A**) SDS-PAGE gel lane of protein sample used for cryo-EM data collection resulting in a symmetry-averaged Sl-Enc1 shell map. (**B**) SDS-PAGE gel lane of protein sample used for cryo-EM data collection resulting in a symmetry-averaged Sl-Enc2 shell map.

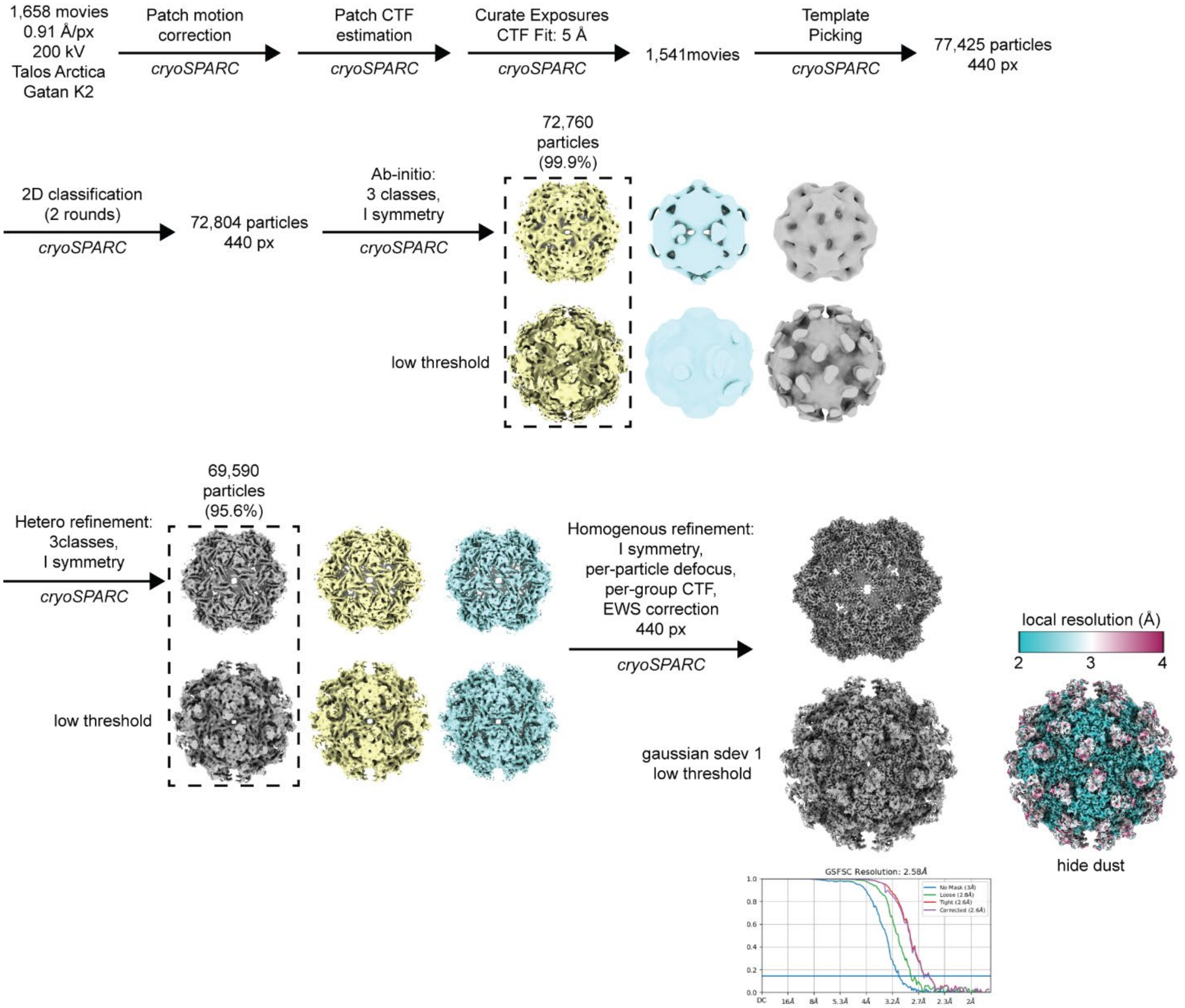


**Fig. S5. Majority Sl-Enc1 cryo-EM data processing workflow.** Cryo-EM workflow showing gold-standard Fourier shell correlation (FSC) curves for the final symmetry-averaged (I) consensus map as well as a local resolution analysis.

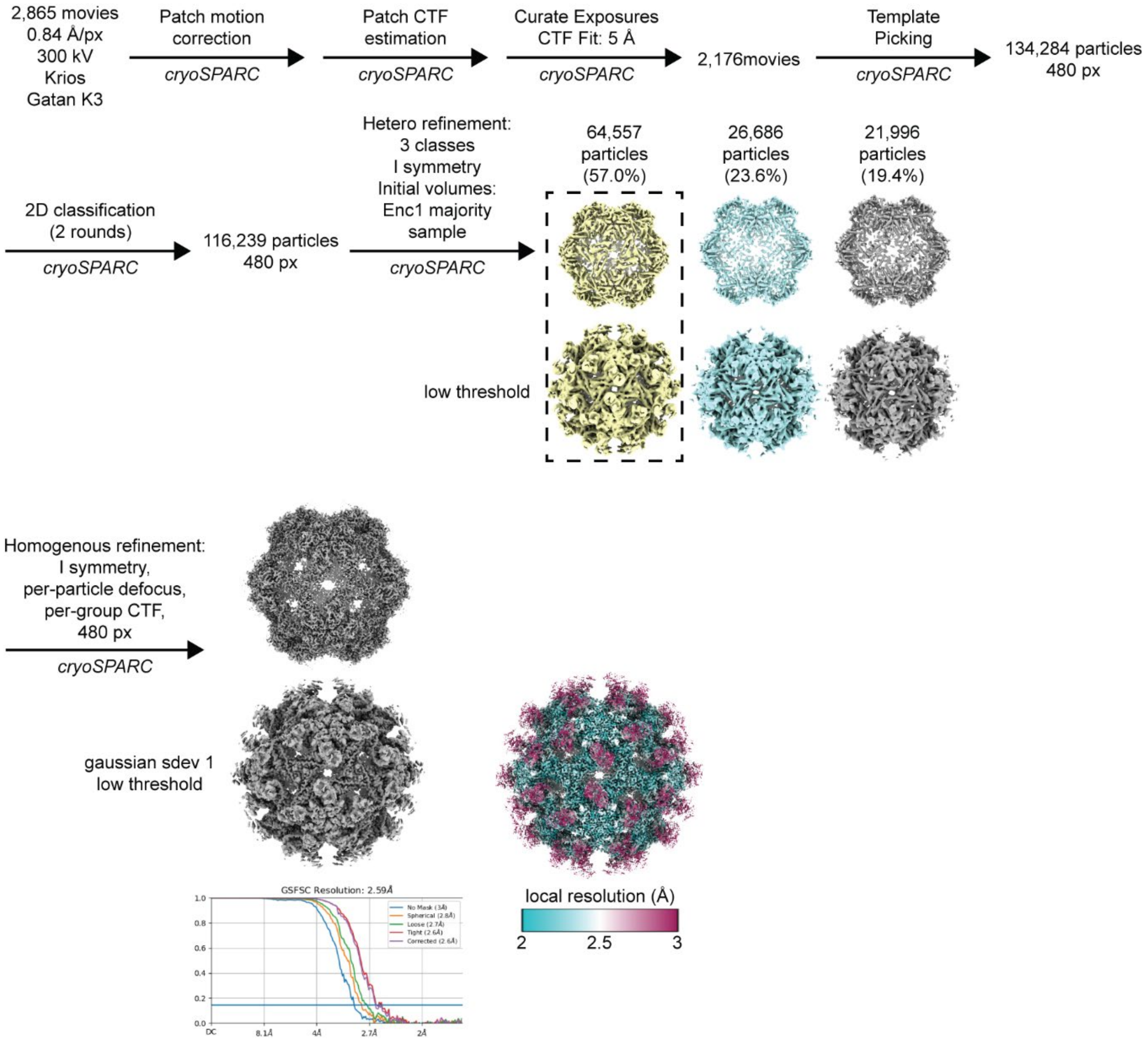


**Fig. S6. Majority SI-Enc2 cryo-EM data processing workflow.** Cryo-EM workflow showing gold-standard Fourier shell correlation (FSC) curves for the final symmetry-averaged (I) consensus map as well as a local resolution analysis.

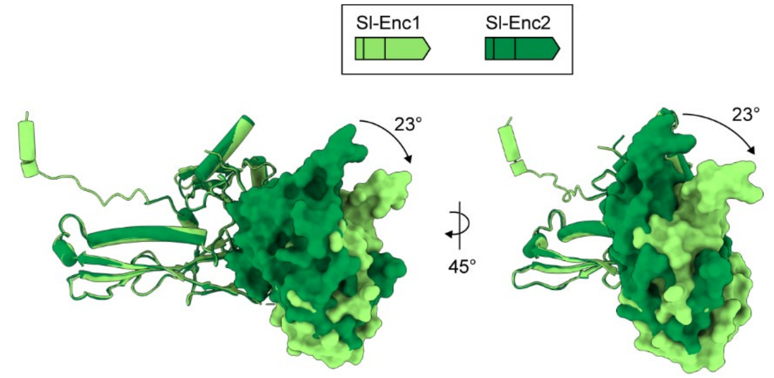


**Fig. S7. Comparison of CBD orientations within Sl-Enc1 and Sl-Enc2.** Structural alignment of the Sl-Enc1 and Sl-Enc2 proteins highlighting the different CBD tilts relative to the HK97-domain. A relative tilt between the Sl-Enc2 and Sl-Enc1 CBDs of ca. 23° can be observed. CBDs shown as surface, HK97-domain as ribbon.

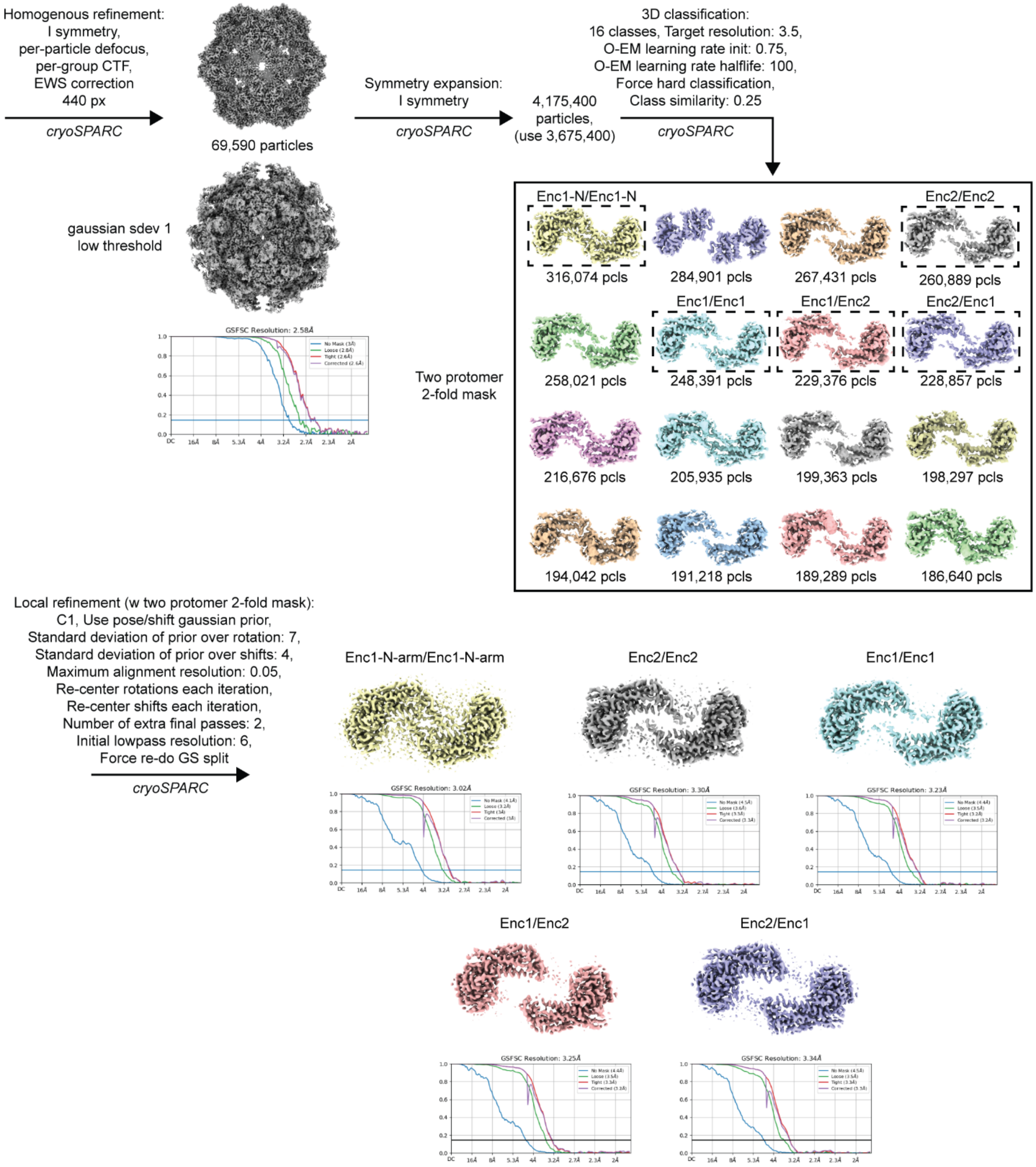


**Fig. S8. Cryo-EM 3D classification analysis workflow for the 2-fold (pore) interaction.**

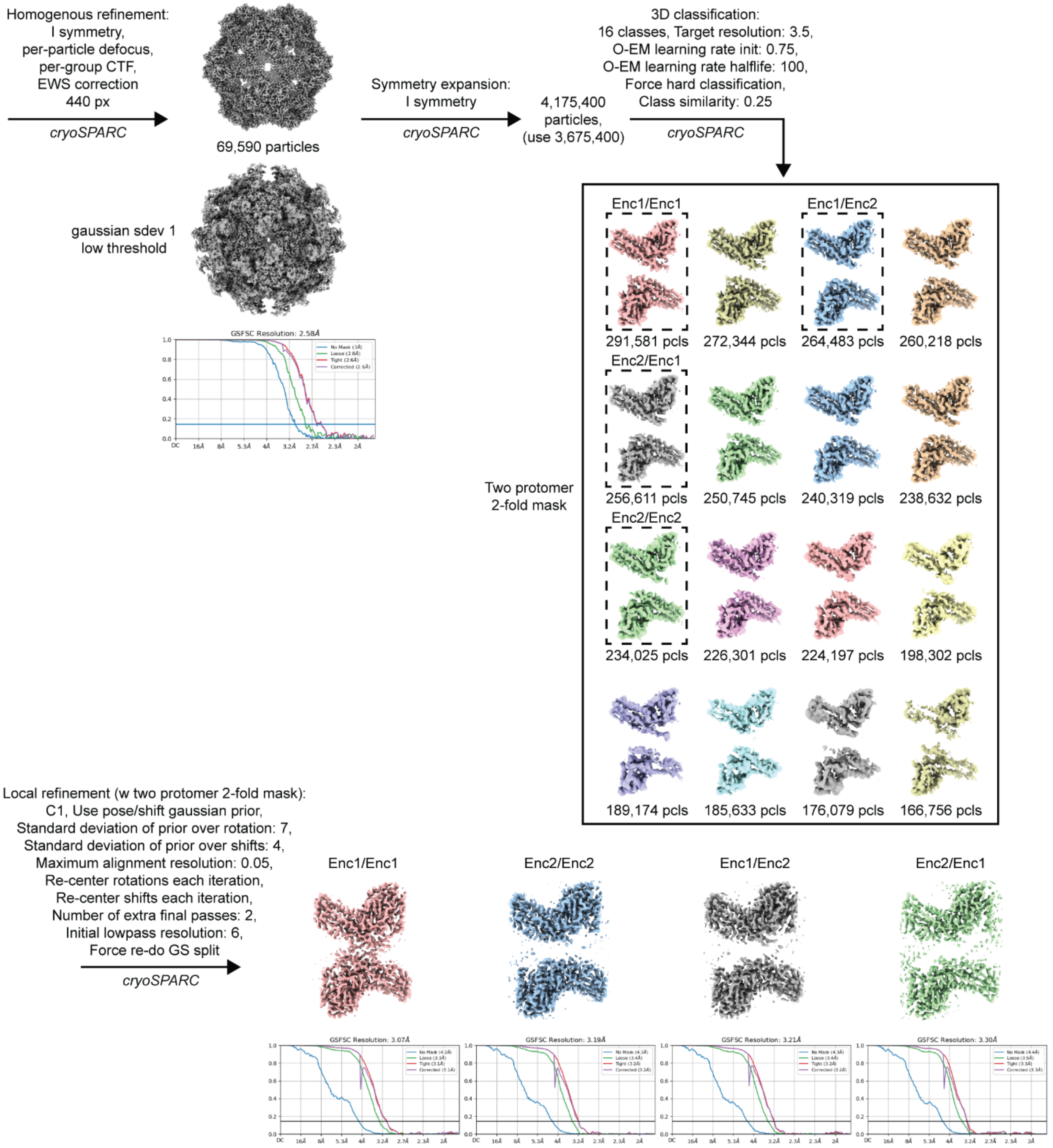


**Fig. S9. Cryo-EM 3D classification analysis workflow for the 2-fold (P-domain) interaction.**

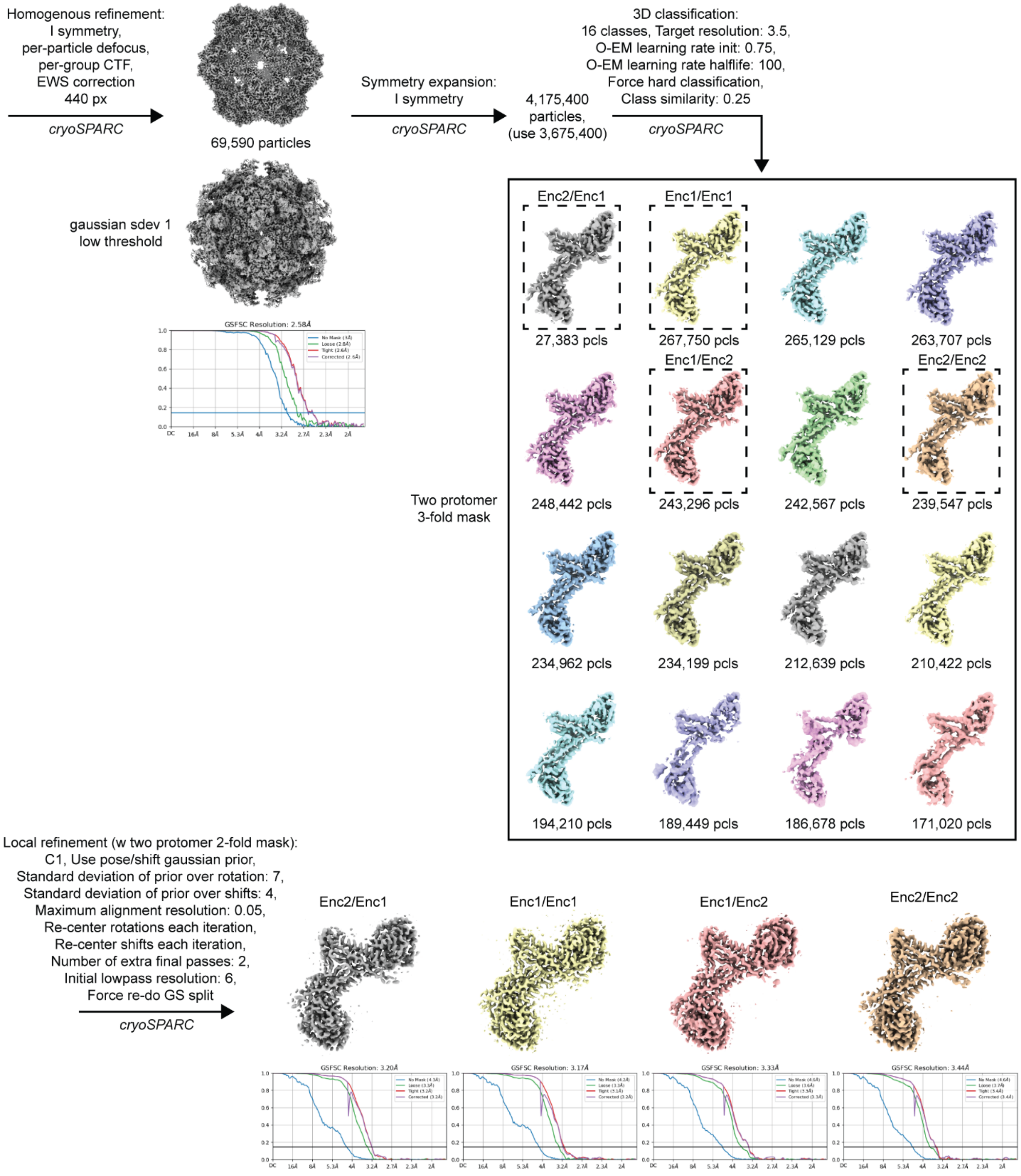


**Fig. S10. Cryo-EM 3D classification analysis workflow for the 3-fold interaction.**

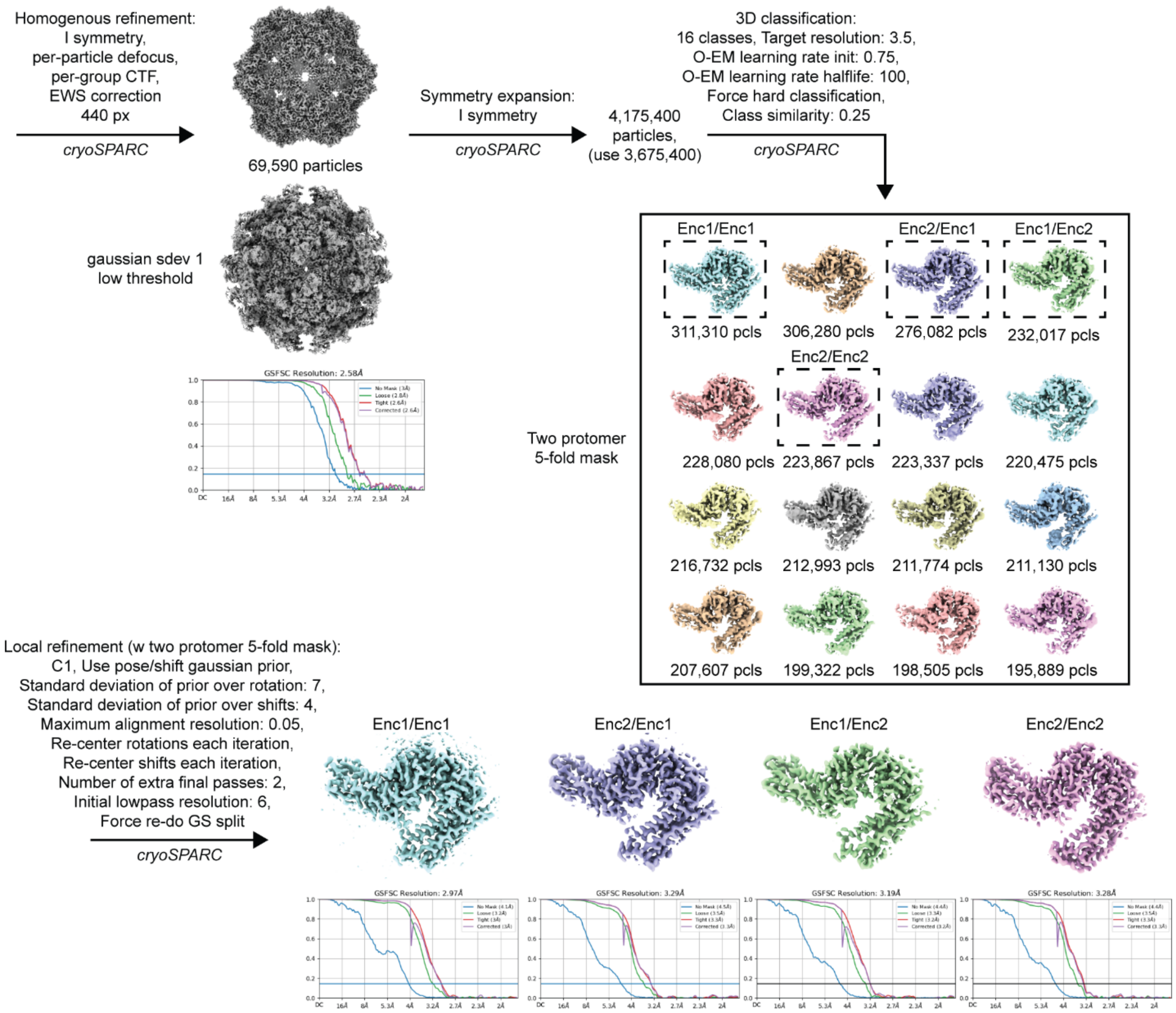


**Fig. S11. Cryo-EM 3D classification analysis workflow for the 5-fold interaction.**

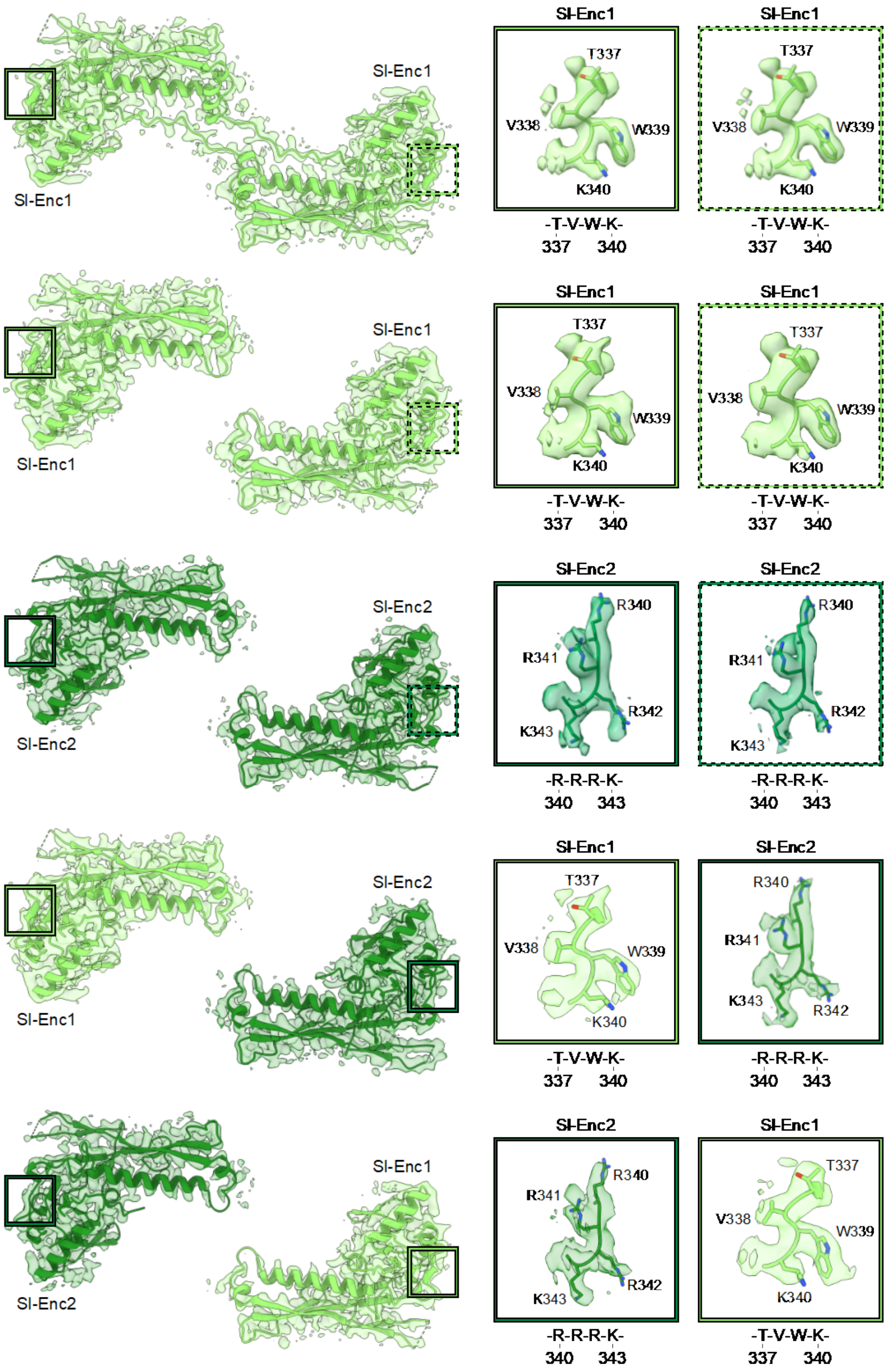


**Fig. S12. Identified high quality classes for the 2-fold (pore) 3D classification analysis.** Left: Cryo-EM densities and atomic models for the five identified classes representing all possible interaction permutations. The class shown on top represents the Sl-Enc1/Sl-Enc1 interaction with resolved N-arm. Right: Diagnostic regions located at the edge of the A-domain used to distinguish between Sl-Enc1 and Sl-Enc2.

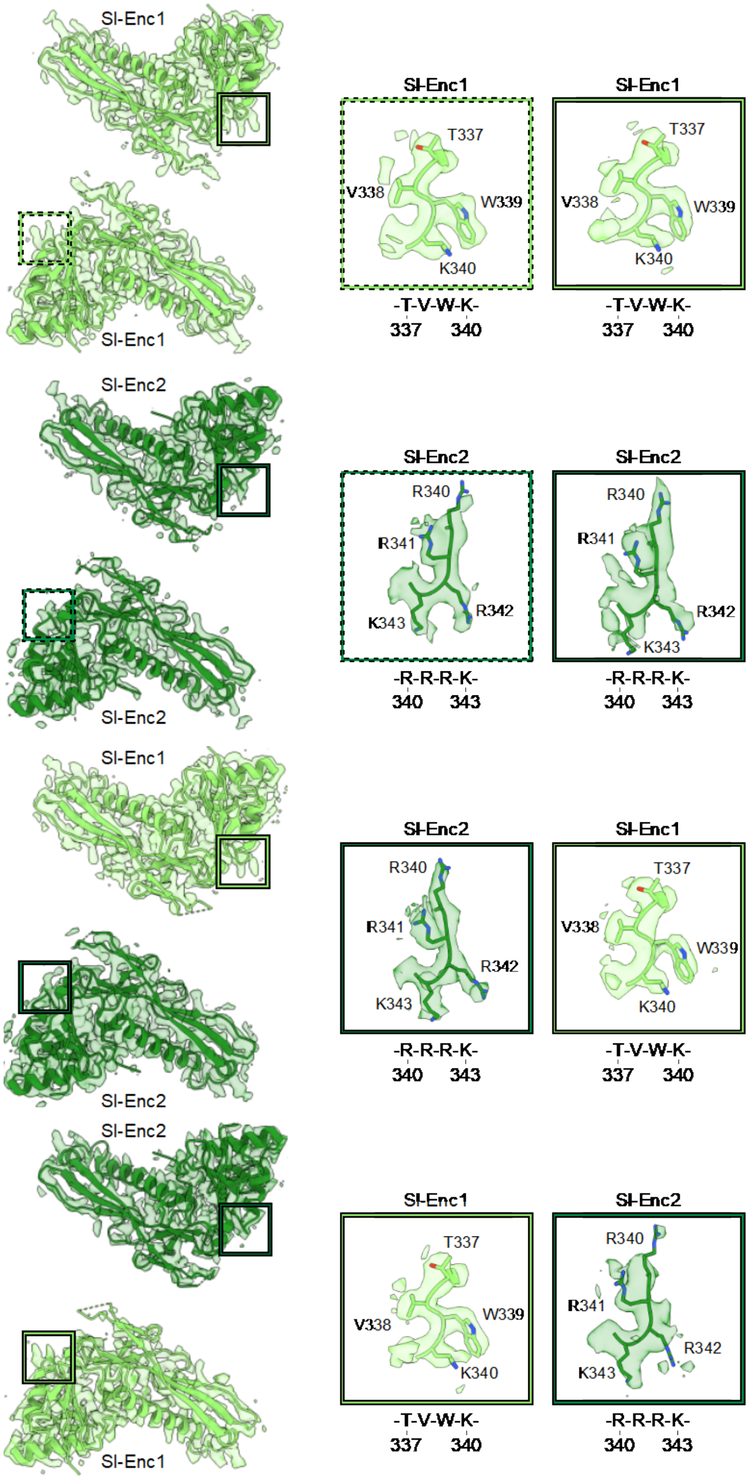


**Fig. S13. Identified high quality classes for the 2-fold (P-domain) 3D classification analysis.** Left: Cryo-EM densities and atomic models for the four identified classes representing all possible interaction permutations. Right: Diagnostic regions located at the edge of the A-domain used to distinguish between Sl-Enc1 and Sl-Enc2.

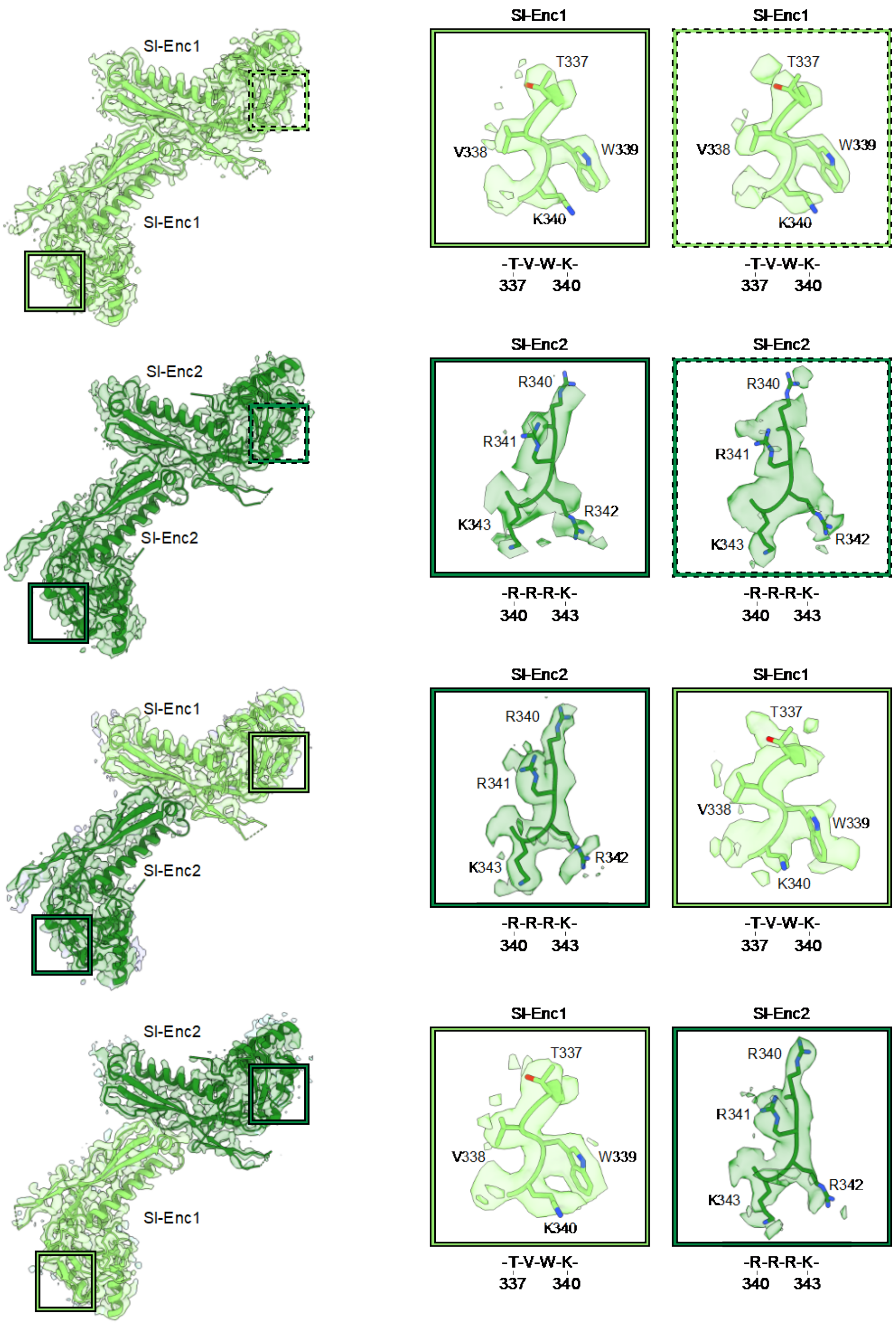


**Fig. S14. Identified high quality classes for the 3-fold 3D classification analysis.** Left: Cryo-EM densities and atomic models for the four identified classes representing all possible interaction permutations. Right: Diagnostic regions located at the edge of the A-domain used to distinguish between SI-Enc1 and SI-Enc2.

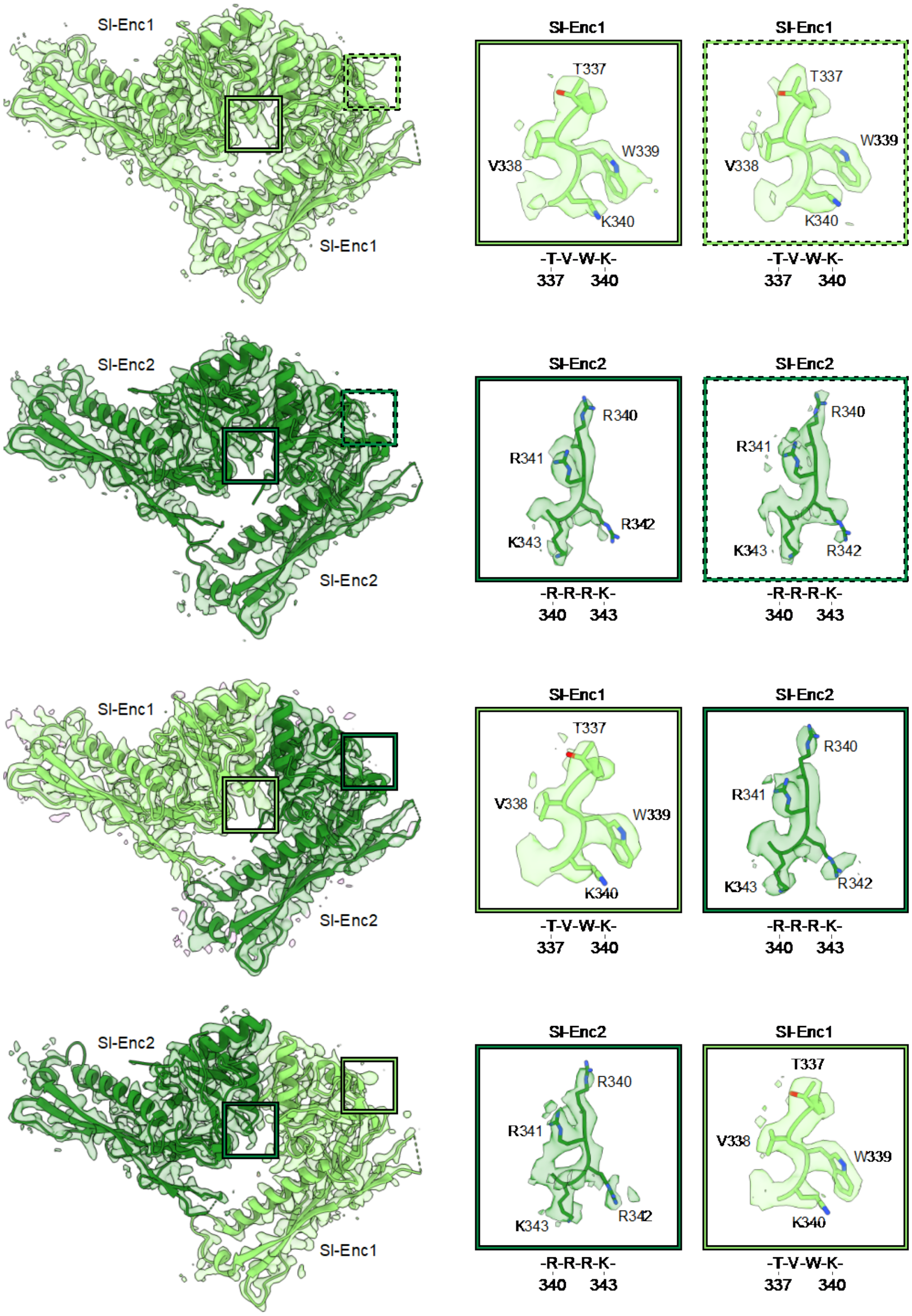


**Fig. S15. Identified high quality classes for the 5-fold 3D classification analysis.** Left: Cryo-EM densities and atomic models for the four identified classes representing all possible interaction permutations. Right: Diagnostic regions located at the edge of the A-domain used to distinguish between Sl-Enc1 and Sl-Enc2.

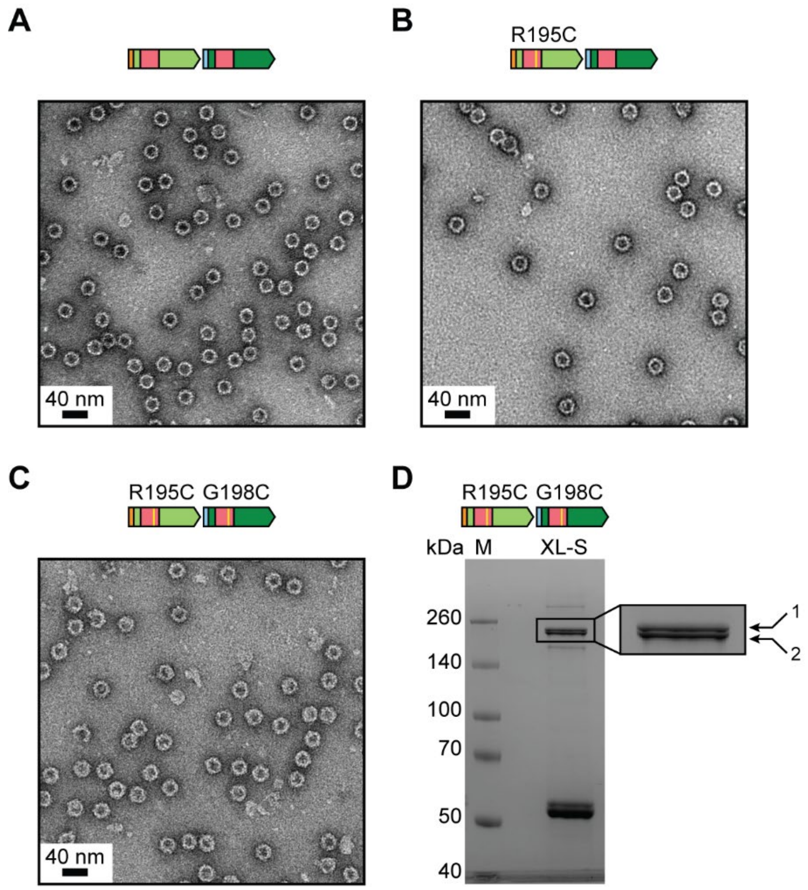


**Fig. S16. Cysteine mutant shell assembly and crosslinking mass spectrometry.** (**A**) Negative stain TEM micrograph of a mixed shell sample containing His-Sl-Enc1 and Strep-Sl-Enc2. (**B**) Negative stain TEM micrograph of a mixed shell sample containing mutant His-Sl-Enc1(R195C) and Strep-Sl-Enc2. (**C**) Negative stain TEM micrograph of a mixed shell sample containing mutant His-Sl-Enc1(R195C) and mutant Strep-Sl-Enc2 (G198C). (**D**) SDS-PAGE analysis of crosslinked double-cysteine mutant sample used for mass spectrometric analysis. M: molecular weight marker. XL-S: Crosslinked His-Sl-Enc1(R195C) + Strep-Sl-Enc2 (G198C) shells.

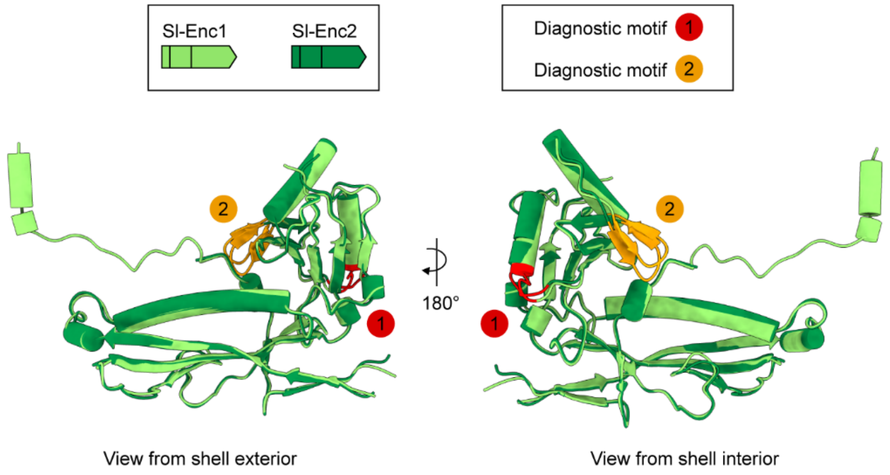


**Fig. S17. Location of the diagnostic motifs within the Sl-Enc1 and Sl-Enc2 protomers.** The two conserved differences between Enc1 and Enc2 shell proteins are highlighted. CBDs not shown for visual clarity.

| gBlock | DNA Sequence |
|---|---|
| Sl-Enc1 | AAGTATAAGAAGGAGATATACAATGACAATCGTAGACGAGACGTTAA<br>ATGGGAACCCAGAGGACACCGCCCATTCGTCTTTGTCTACGGCTGC<br>CGCCCGCAACTTGGCCACCACAACCAAAACTGTACCGCAAATGCAG<br>GGCATCACATCCCGTTGGTTGCTTCGTTTGTTACCGTGGGTTCAGG<br>TCTCAGGGGGCACGTACCGCGTAAACCGCCGTTTATCACACACCGT<br>GGGGGATGGACGTATCGACTTTGATATTTCTGGTAGTGACGTAGCG<br>ATTATCCCCGAAGAATTACGCGAACTTCCAGCGCTTCGCGACTTCA<br>CGGATACTGAGGTCCTTGCTGCATTGGGGGAGCGTTTCACTCAGCG<br>TGAGTATGCCCCAGGAGAGTTGATCGCAGAGGCTGGGCGCCCAGG<br>GGATCGCCTTGTTTTAATCGCGCATGGCCGCGTAGATCGCATTGGA<br>ACTGGAAAGTACGGCGACACCACCGTACTTGCAGCGCTTGCGGGG<br>GGCGACCACCTGGGGGACGCGCCGCTTACCTCTGACGAAGTTACA<br>TGGGAATTTTCGTATCGCGCGGTTACTCGCGTAACAGTAGTCGAGC<br>TTCCGCGCCGCGCCGCCTTAGAAATCATCGAACGTTCCCCAGGTCT<br>TCGTGAGCATTTAGCCGGGGTACGTCAAGGGCCGGTTCACCCCAC<br>AAACAGTTCCGGCGAATCTGCTGTAGCCGTGGCTAGTGGTCACCGT<br>GGCGAGCCGTCGCTGCCCGGAGCGTTCGCGGATTATGACTTAGCG<br>CCCCGCGAGTACGAACTTAGCGTCGCTCAGACAGTATTGCGCACGC<br>ATACGCGCGTGGGGGATTTATACAATGATCCTATGAACCAGGTGGA<br>AGAACAATTAAAATTGACCGTCCAGGCACTGCGCGAACGCCAGGAA<br>CATGAGATGATCAACAATCGTGAGTTCGGATTGCTTCACAATGCAGA<br>CTTAAAGCAACGTATTCCTACGCGTAGCGGACCGCCTACTCCTGAT<br>GACTTGGATGATTTGCTGGCGACAGTTTGGAAGGACCCTGGCTTTC<br>TTTTAGCGCATCCACGCGCCATCGCAGCAATGGCACGTGAATGGTC<br>GGCTCGCGGACTGTATCCAACGGCGGTCGATTTTCATGGGCATTCT<br>CTTCCATCCTGGCGTGGTGTTCCAATCTTTCCGTGCAATAAAATTCC<br>AGTCACTAAGGAACGCACAAGCTCCATTCTGCTTCTTCGCACCGGC<br>GAGGAAAAGCAGGGCGTCGTCGGCCTTCACCAAACTGGGATTCCT<br>GACGAGTATGAGCCAAGTCTGTCAGTTCGTTTCATGGGAATCGATG<br>ACCGCGCTGTAATTAACTACTTGGTTTCTGCGTACTACAGTGCAGCC<br>GTGCTTGTACCAGATGCATTGGGAGTCTTAGAGGATGTCGAAGTCG<br>GGTTATGAATTAACCTAGGCTGCTGC |

| | |
|---|---|
| SI-Enc2 | AAGTATAAGAAGGAGATATACAATGACTACGTCCGTGGATCCAACTT<br>CAGGTCCTCAGGCAAGTGGGACAGAGCAGAATCGCTCAAGCTTGG<br>ACACAGCCGCTGCTCGTAAACTGGCTACTACCACGAAAACGGTCCC<br>GCAAATGCAGGGCATTTCATCACGCTGGTTGCTTCGTGTCTTACCG<br>TGGACGCAGGTCAATGGGGGTACATACCGTGTTAATCGCCGCCTGA<br>CTCATACCCTGGGAGACGGTCAAGTGGAGTTCGTAAACACCGGCG<br>CTGAAGTTCGTGTAATTCCGGAGGAGCTGCGTGAGTTGGCCCCGCT<br>TCGTGGGTTTACCGATACTCCAACTCTGGAAGCCCTTGCGGGTCGC<br>TTTGCACAACACGAGTTTGCTCCAGGCGATGTCTTAGCTCAGCAGG<br>ACCAACCGGCTGACCGCATCATCTTGATTGCTCACGGTAAGTTGGA<br>TCGTTTAGGCACTGGCAAGTATGGCGGAGAAACGGTGCAAGGGCA<br>GTTGGCCGGCGGTGACCACCTGGGAGCGGCCGCACTGTTAGACGG<br>TGGTGCGGCCTGGGAGCATACTGTCCGCGCTGTTACCCGCGTCAC<br>TGCACTGACTTTATCGCGCGGTGATTATGAACAAGTCCTGGGTGGT<br>TCCGAGAGCTTACGTGCGCACGTCGAAGCTTTTCGTGCGGCCTTAG<br>TACCGGCTCAGAATAAGCACGGCGAGGCTGCGATCGAAGTAGCTG<br>CCGGTCATGTCGGGGAACCCTCCCTGCCAGGGACGTTTGCCGATT<br>ACGATCTTGCTCCTCGTGAGTACGAGCTTTCAGTAGCCCAGACGGT<br>TTTAAAGATTCACTCTCGTGTTGCCGACCTGTATAATGACCCCATGA<br>ACCAAATGGATCAGCAATTACGCTTAACTGTCGAGGCTTTGCGCGA<br>ACGCCAAGAACATGAGATGATCAACAACCGTGAATTTGGTTTACTGC<br>ATAATGCAGATTTAAAGCAGCGCATCCATACGCGCTCCGGCCCCCC<br>AACACCTGACGACCTGGACGAGTTAATCTCTCGCCGCCGCAAAACG<br>CAAGTGTTGTTGGCGCATCCACGCACTATTGCCGCCATTGGTCGTG<br>AGTGGAACGCGCGTGGAATCTATCCGACGGGCGCCGAATTGCATG<br>GAACAGATGTACGCGCTTGGCGTGGAATTCCTTTACTTCCTTGTAAC<br>AAAATTCCCGTAACCCCGGAACAAACAAGCAGTATCATTGCTATGCG<br>CTTGGGTGAGGAAAACCAGGGGGTAGTCGGATTGCACCAGACCGG<br>CATTCCTGACGAGTATCAACCAGGCTTGTCTGTTCGTTTTATGGGGA<br>TTAATGATCAGGCAGTCATCCAGTATCTTGTTAGCGCGTACTATTCA<br>GCAGCAGTGTTGGTGCCTGATGCTTTGGGGATTCTGGAGGATGTTG<br>AAATCGGGCATTGAATTAACCTAGGCTGCTGC |

| | |
|---|---|
| SI-Enc1+SI-Enc2 | AAGTATAAGAAGGAGATATACAATGACAATCGTAGACGAGACGTTAA<br>ATGGGAACCCAGAGGACACCGCCCATTCGTCTTTGTCTACGGCTGC<br>CGCCCGCAACTTGGCCACCACAACCAAAACTGTACCGCAAATGCAG<br>GGCATCACATCCCGTTGGTTGCTTCGTTTGTTACCGTGGGTTCAGG<br>TCTCAGGGGGCACGTACCGCGTAAACCGCCGTTTATCACACACCGT<br>GGGGGATGGACGTATCGACTTTGATATTTCTGGTAGTGACGTAGCG<br>ATTATCCCCGAAGAATTACGCGAACTTCCAGCGCTTCGCGACTTCA<br>CGGATACTGAGGTCCTTGCTGCATTGGGGGAGCGTTTCACTCAGCG<br>TGAGTATGCCCCAGGAGAGTTGATCGCAGAGGCTGGGCGCCCAGG<br>GGATCGCCTTGTTTTAATCGCGCATGGCCGCGTAGATCGCATTGGA<br>ACTGGAAAGTACGGCGACACCACCGTACTTGCAGCGCTTGCGGGG<br>GGCGACCACCTGGGGGACGCGCCGCTTACCTCTGACGAAGTTACA<br>TGGGAATTTTCGTATCGCGCGGTTACTCGCGTAACAGTAGTCGAGC<br>TTCCGCGCCGCGCCGCCTTAGAAATCATCGAACGTTCCCCAGGTCT<br>TCGTGAGCATTTAGCCGGGGTACGTCAAGGGCCGGTTCACCCCAC<br>AAACAGTTCCGGCGAATCTGCTGTAGCCGTGGCTAGTGGTCACCGT<br>GGCGAGCCGTCGCTGCCCGGAGCGTTCGCGGATTATGACTTAGCG<br>CCCCGCGAGTACGAACTTAGCGTCGCTCAGACAGTATTGCGCACGC<br>ATACGCGCGTGGGGGATTTATACAATGATCCTATGAACCAGGTGGA<br>AGAACAATTAAAATTGACCGTCCAGGCACTGCGCGAACGCCAGGAA<br>CATGAGATGATCAACAATCGTGAGTTCGGATTGCTTCACAATGCAGA<br>CTTAAAGCAACGTATTCCTACGCGTAGCGGACCGCCTACTCCTGAT<br>GACTTGGATGATTTGCTGGCGACAGTTTGGAAGGACCCTGGCTTTC<br>TTTTAGCGCATCCACGCGCCATCGCAGCAATGGCACGTGAATGGTC<br>GGCTCGCGGACTGTATCCAACGGCGGTCGATTTTCATGGGCATTCT<br>CTTCCATCCTGGCGTGGTGTTCCAATCTTTCCGTGCAATAAAATTCC<br>AGTCACTAAGGAACGCACAAGCTCCATTCTGCTTCTTCGCACCGGC<br>GAGGAAAAGCAGGGCGTCGTCGGCCTTCACCAAACTGGGATTCCT<br>GACGAGTATGAGCCAAGTCTGTCAGTTCGTTTCATGGGAATCGATG<br>ACCGCGCTGTAATTAACTACTTGGTTTCTGCGTACTACAGTGCAGCC<br>GTGCTTGTACCAGATGCATTGGGAGTCTTAGAGGATGTCGAAGTCG<br>GGTTATGATATAAGAAGGAGATATACATGACTACGTCCGTGGATCCA<br>ACTTCAGGTCCTCAGGCAAGTGGGACAGAGCAGAATCGCTCAAGCT<br>TGGACACAGCCGCTGCTCGTAAACTGGCTACTACCACGAAAACGGT<br>CCCGCAAATGCAGGGCATTTCATCACGCTGGTTGCTTCGTGTCTTA<br>CCGTGGACGCAGGTCAATGGGGGTACATACCGTGTTAATCGCCGC<br>CTGACTCATACCCTGGGAGACGGTCAAGTGGAGTTCGTAAACACCG<br>GCGCTGAAGTTCGTGTAATTCCGGAGGAGCTGCGTGAGTTGGCCC<br>CGCTTCGTGGGTTTACCGATACTCCAACTCTGGAAGCCCTTGCGGG<br>TCGCTTTGCACAACACGAGTTTGCTCCAGGCGATGTCTTAGCTCAG<br>CAGGACCAACCGGCTGACCGCATCATCTTGATTGCTCACGGTAAGT<br>TGGATCGTTTAGGCACTGGCAAGTATGGCGGAGAAACGGTGCAAG<br>GGCAGTTGGCCGGCGGTGACCACCTGGGAGCGGCCGCACTGTTAG<br>ACGGTGGTGCGGCCTGGGAGCATACTGTCCGCGCTGTTACCCGCG<br>TCACTGCACTGACTTTATCGCGCGGTGATTATGAACAAGTCCTGGG<br>TGGTTCCGAGAGCTTACGTGCGCACGTCGAAGCTTTTCGTGCGGCC |

| | |
|---|---|
| | TTAGTACCGGCTCAGAATAAGCACGGCGAGGCTGCGATCGAAGTAG<br>CTGCCGGTCATGTCGGGGAACCCTCCCTGCCAGGGACGTTTGCCG<br>ATTACGATCTTGCTCCTCGTGAGTACGAGCTTTCAGTAGCCCAGAC<br>GGTTTTAAAGATTCACTCTCGTGTTGCCGACCTGTATAATGACCCCA<br>TGAACCAAATGGATCAGCAATTACGCTTAACTGTCGAGGCTTTGCG<br>CGAACGCCAAGAACATGAGATGATCAACAACCGTGAATTTGGTTTAC<br>TGCATAATGCAGATTTAAAGCAGCGCATCCATACGCGCTCCGGCCC<br>CCCAACACCTGACGACCTGGACGAGTTAATCTCTCGCCGCCGCAAA<br>ACGCAAGTGTTGTTGGCGCATCCACGCACTATTGCCGCCATTGGTC<br>GTGAGTGGAACGCGCGTGGAATCTATCCGACGGGCGCCGAATTGC<br>ATGGAACAGATGTACGCGCTTGGCGTGGAATTCCTTTACTTCCTTGT<br>AACAAAATTCCCGTAACCCCGGAACAAACAAGCAGTATCATTGCTAT<br>GCGCTTGGGTGAGGAAAACCAGGGGGTAGTCGGATTGCACCAGAC<br>CGGCATTCCTGACGAGTATCAACCAGGCTTGTCTGTTCGTTTTATGG<br>GGATTAATGATCAGGCAGTCATCCAGTATCTTGTTAGCGCGTACTAT<br>TCAGCAGCAGTGTTGGTGCCTGATGCTTTGGGGATTCTGGAGGATG<br>TTGAAATCGGGCATTGATTAACCTAGGCTGCTGC |

**Table S1. DNA sequences of gBlock Gene Fragments.**

| Primer | DNA Sequence |
|---|---|
| His-SI-Enc1-iPCR-FW | ATGCACCACCACCATCACCATACAATCGTAGACGAGACGTTAAATGGG |
| pETDuet-1-iPCR-RV | TGTATATCTCCTTCTTATACTTAACTAATATACTAAGATGGGG |
| His-SI-Enc2-iPCR-FW | ATGCACCACCACCATCACCATACTACGTCCGTGGATCCAACTTC |
| Double-Enc-His-SI-Enc2-iPCR-FW | ACTACGTCCGTGGATCCAACTTC |
| Double-Enc-His-SI-Enc2-iPCR-RV | ATGGTGATGGTGGTGGTGCATGTATATCTCCTTCTTATATCATAACCCG |
| Double-Enc-Strep-SI-Enc2-iPCR-RV | CTTTTCGAACTGCGGGTGGCTCCACATGTATATCTCCTTCTTATATCATAACCC |
| His-SI-Enc1-R195C-FW | TGTGCCGCCTTAGAAATCATCG |
| His-SI-Enc1-R195C-iPCR-RV | GCGCGGAAGCTCGACTAC |
| His-SI-Enc1-R195C-Gibson-Insert-RV | ACAGCGCGATAAAGTCAGTGC |
| His-SI-Enc1- | ACTGACTTTATCGCGCTGTGATTATGAACAAGTCCTGG |

| | |
|---|---|
| R195C-Gibson-Vector-FW | |
| His-SI-Enc1-R195C-Gibson-Vector-RV | GATTTCTAAGGCGGCACAGCGCGGAAGCTCGAC |

**Table S2. DNA sequences of PCR primers used to construct plasmids.**

| **Plasmid** | **Cloning Method** | **Primers** |
|---|---|---|
| His-Sl-Enc1 | Inverse PCR | His-Sl-Enc1-iPCR-FW<br>pETDuet-1-iPCR-RV |
| His-Sl-Enc2 | Inverse PCR | His-Sl-Enc2-iPCR-FW<br>pETDuet-1-iPCR-RV |
| Sl-Enc1 / His-Sl-Enc2 | Inverse PCR | Double-Enc-His-Sl-Enc2-iPCR-FW<br>Double-Enc-His-Sl-Enc2-iPCR-RV |
| His-Sl-Enc1 / Sl-Enc2 | Inverse PCR | His-Sl-Enc1-iPCR-FW<br>pETDuet-1-iPCR-RV |
| His-Sl-Enc1 / His-Sl-Enc2 | Inverse PCR | His-Sl-Enc1-iPCR-FW<br>pETDuet-1-iPCR-RV |
| His-Sl-Enc1 / Strep-Sl-Enc2 | Inverse PCR | Double-Enc-His-Sl-Enc2-iPCR-FW<br>Double-Enc-Strep-Sl-Enc2-iPCR-RV |
| His-Sl-Enc1(R195C) / Strep-Sl-Enc2 | Inverse PCR | His-Sl-Enc1-R195C-FW<br>His-Sl-Enc1-R195C-iPCR-RV |
| His-Sl-Enc1(R195C) / Strep-Sl-Enc2(G198C) | Gibson Assembly | Insert<br>His-Sl-Enc1-R195C-FW<br>His-Sl-Enc1-R195C-Gibson-Insert-RV |
| | | Vector<br>His-Sl-Enc1-R195C-Gibson-Vector-FW<br>His-Sl-Enc1-R195C-Gibson-Vector-RV |

**Table S3. Primer pairs used to construct the plasmids used in this study.**

| Protein | Protein Sequence |
|---|---|
| Sl-Enc1 | MTIVDETLNGNPEDTAHSSLSTAAARNLATTTKTVPQMQGITSRWLLRL LPWVQVSGGTYRVNRRLSHTVGDGRIDFDISGSDVAIIPEELRELPALR DFTDTEVLAALGERFTQREYAPGELIAEAGRPGDRLVLIAHGRVDRIGT GKYGDTTVLAALAGGDHLGDAPLTSDEVTWEFSYRAVTRVTVVELPRR AALEIIERSPGLREHLAGVRQGPVHPTNSSGESAVAVASGHRGEPSLP GAFADYDLAPREYELSVAQTVLRTHTRVGDLYNDPMNQVEEQLKLTVQ ALRERQEHEMINNREFGLLHNADLKQRIPTRSGPPTPDDLDDLLATVW KDPGFLLAHPRAIAAMAREWSARGLYPTAVDFHGHSLPSWRGVPIFPC NKIPVTKERTSSILLLRTGEEKQGVVGLHQTGIPDEYEPSLSVRFMGIDD RAVINYLVSAYYSAAVLVPDALGVLEDVEVGL* |
| His-Sl-Enc1 | MHHHHHHTIVDETLNGNPEDTAHSSLSTAAARNLATTTKTVPQMQGIT SRWLLRLLPWVQVSGGTYRVNRRLSHTVGDGRIDFDISGSDVAIIPEEL RELPALRDFTDTEVLAALGERFTQREYAPGELIAEAGRPGDRLVLIAHG RVDRIGTGKYGDTTVLAALAGGDHLGDAPLTSDEVTWEFSYRAVTRVT VVELPRRAALEIIERSPGLREHLAGVRQGPVHPTNSSGESAVAVASGH RGEPSLPGAFADYDLAPREYELSVAQTVLRTHTRVGDLYNDPMNQVE EQLKLTVQALRERQEHEMINNREFGLLHNADLKQRIPTRSGPPTPDDL DDLLATVWKDPGFLLAHPRAIAAMAREWSARGLYPTAVDFHGHSLPS WRGVPIFPCNKIPVTKERTSSILLLRTGEEKQGVVGLHQTGIPDEYEPSL SVRFMGIDDRAVINYLVSAYYSAAVLVPDALGVLEDVEVGL* |
| His-Sl-Enc1 (R195C) | MHHHHHHTIVDETLNGNPEDTAHSSLSTAAARNLATTTKTVPQMQGIT SRWLLRLLPWVQVSGGTYRVNRRLSHTVGDGRIDFDISGSDVAIIPEEL RELPALRDFTDTEVLAALGERFTQREYAPGELIAEAGRPGDRLVLIAHG RVDRIGTGKYGDTTVLAALAGGDHLGDAPLTSDEVTWEFSYRAVTRVT VVELPRCAALEIIERSPGLREHLAGVRQGPVHPTNSSGESAVAVASGH RGEPSLPGAFADYDLAPREYELSVAQTVLRTHTRVGDLYNDPMNQVE EQLKLTVQALRERQEHEMINNREFGLLHNADLKQRIPTRSGPPTPDDL DDLLATVWKDPGFLLAHPRAIAAMAREWSARGLYPTAVDFHGHSLPS WRGVPIFPCNKIPVTKERTSSILLLRTGEEKQGVVGLHQTGIPDEYEPSL SVRFMGIDDRAVINYLVSAYYSAAVLVPDALGVLEDVEVGL* |
| Sl-Enc2 | MTTSVDPTSGPQASGTEQNRSSLDTAAARKLATTTKTVPQMQGISSR WLLRVLPWTQVNGGTYRVNRRLTHTLGDGQVEFVNTGAEVRVIPEEL RELAPLRGFTDTPTLEALAGRFAQHEFAPGDVLAQQDQPADRIILIAHG KLDRLGTGKYGGETVQGQLAGGDHLGAAALLDGGAAWEHTVRAVTR VTALTLSRGDYEQVLGGSESLRAHVEAFRAALVPAQNKHGEAAIEVAA GHVGEPSLPGTFADYDLAPREYELSVAQTVLKIHSRVADLYNDPMNQM DQQLRLTVEALRERQEHEMINNREFGLLHNADLKQRIHTRSGPPTPDD LDELISRRRKTQVLLAHPRTIAAIGREWNARGIYPTGAELHGTDVRAWR GIPLLPCNKIPVTPEQTSSIIAMRLGEENQGVVGLHQTGIPDEYQPGLSV RFMGINDQAVIQYLVSAYYSAAVLVPDALGILEDVEIGH* |

| | |
|---|---|
| His-SI-Enc2 | MHHHHHHTTSVDPTSGPQASGTEQNRSSLDTAAARKLATTTKTVPQMQGISSRWLLRVLPWTQVNGGTYRVNRRLTHTLGDGQVEFVNTGAEVRVIPEELRELAPLRGFTDTPTLEALAGRFAQHEFAPGDVLAQQDQPADRIILIAHGKLDRLGTGKYGGETVQGQLAGGDHLGAAALLDGGAAWEHTVRAVTRVTALTLSRGDYEQVLGGSESLRAHVEAFRAALVPAQNKHGEAAIEVAAGHVGEPSLPGTFADYDLAPREYELSVAQTVLKIHSRVADLYNDPMNQMDQQLRLTVEALRERQEHEMINNREFGLLHNADLKQRIHTRSGPPTPDDLDELISRRRKTQVLLAHPRTIAAIGREWNARGIYPTGAELHGTDVRAWRGIPLLPCNKIPVTPEQTSSIIAMRLGEENQGVVGLHQTGIPDEYQPGLSVRFMGINDQAVIQYLVSAYYSAAVLVPDALGILEDVEIGH* |
| Strep-SI-Enc2 | MWSHPQFEKTTSVDPTSGPQASGTEQNRSSLDTAAARKLATTTKTVPQMQGISSRWLLRVLPWTQVNGGTYRVNRRLTHTLGDGQVEFVNTGAEVRVIPEELRELAPLRGFTDTPTLEALAGRFAQHEFAPGDVLAQQDQPADRIILIAHGKLDRLGTGKYGGETVQGQLAGGDHLGAAALLDGGAAWEHTVRAVTRVTALTLSRGDYEQVLGGSESLRAHVEAFRAALVPAQNKHGEAAIEVAAGHVGEPSLPGTFADYDLAPREYELSVAQTVLKIHSRVADLYNDPMNQMDQQLRLTVEALRERQEHEMINNREFGLLHNADLKQRIHTRSGPPTPDDLDELISRRRKTQVLLAHPRTIAAIGREWNARGIYPTGAELHGTDVRAWRGIPLLPCNKIPVTPEQTSSIIAMRLGEENQGVVGLHQTGIPDEYQPGLSVRFMGINDQAVIQYLVSAYYSAAVLVPDALGILEDVEIGH* |
| Strep-SI-Enc2 (G198C) | MWSHPQFEKTTSVDPTSGPQASGTEQNRSSLDTAAARKLATTTKTVPQMQGISSRWLLRVLPWTQVNGGTYRVNRRLTHTLGDGQVEFVNTGAEVRVIPEELRELAPLRGFTDTPTLEALAGRFAQHEFAPGDVLAQQDQPADRIILIAHGKLDRLGTGKYGGETVQGQLAGGDHLGAAALLDGGAAWEHTVRAVTRVTALTLSRCDYEQVLGGSESLRAHVEAFRAALVPAQNKHGEAAIEVAAGHVGEPSLPGTFADYDLAPREYELSVAQTVLKIHSRVADLYNDPMNQMDQQLRLTVEALRERQEHEMINNREFGLLHNADLKQRIHTRSGPPTPDDLDELISRRRKTQVLLAHPRTIAAIGREWNARGIYPTGAELHGTDVRAWRGIPLLPCNKIPVTPEQTSSIIAMRLGEENQGVVGLHQTGIPDEYQPGLSVRFMGINDQAVIQYLVSAYYSAAVLVPDALGILEDVEIGH* |

**Table S4. Protein sequences used in this work.**

| | Sl-Enc1 (EMD-44632) (PDB ID: 9BJE) | Sl-Enc2 (EMD-44603) (PDB ID: 9BIX) |
|---|---|---|
| **Data collection and processing** | | |
| Magnification | 45,000x | 105,000x |
| Voltage (kV) | 200 | 300 |
| Electron exposure (e–/Å$^2$) | 45.50 | 54.0 |
| Defocus range (μm) | -0.8 to -1.8 | -0.5 to -1.0 |
| Pixel size (Å) | 0.91 | 0.84 |
| Symmetry imposed | I | I |
| Initial particle images (no.) | 77,425 | 288,651 |
| Final particle images (no.) | 69,590 | 155,190 |
| Map resolution (Å)<br>FSC threshold | 2.58<br>0.143 | 2.59<br>0.143 |
| **Refinement** | | |
| Initial model used (PDB code) | 9BIX | 9BHU |
| Model resolution (Å)<br>FSC threshold | 2.8<br>0.5 | 2.7<br>0.5 |
| Map sharpening *B* factor (Å$^2$) | -99.9 | -114.4 |
| Model composition<br>Non-hydrogen atoms<br>Protein residues<br>Ligands | <br>2,094<br>266<br>0 | <br>1,944<br>244<br>0 |
| *B* factors (Å$^2$)<br>Protein<br>Ligands | <br>39.30<br>- | <br>33.63<br>- |
| R.m.s. deviations<br>Bond lengths (Å)<br>Bond angles (°) | <br>0.005<br>1.112 | <br>0.003<br>0.513 |
| Validation<br>MolProbity score<br>Clashscore<br>Poor rotamers (%) | <br>1.41<br>4.53<br>0.88 | <br>1.44<br>3.34<br>0.95 |
| Ramachandran plot<br>Favored (%)<br>Allowed (%)<br>Disallowed (%) | <br>96.95<br>3.05<br>0 | <br>95.42<br>4.58<br>0 |

**Table S5. Cryo-EM data collection, refinement, and validation statistics.**
